# Endosomal mRNA transport coordinates local mitochondrial bioenergetics during polar fungal growth

**DOI:** 10.64898/2026.06.12.731371

**Authors:** Johannes Postma, Pascal Künzel, Simon Wegmann, Kira Müntjes, Senthil Kumar Devan, Srimeenakshi Sankaranarayanan, Stephan Krueger, Philipp Westhoff, Nick Wierckx, Lasse van Wijlick, Michael Feldbrügge

## Abstract

Mitochondrial function relies on the precise spatial coordination of protein synthesis and import. Most mitochondrial proteins are nuclear-encoded and must be supplied across varying intracellular distances. In highly polarized cells such as fungal hyphae and neurons, active long-distance mRNA transport is thought to sustain distal mitochondrial function, but its mechanistic coupling to protein import and organelle physiology is unclear. Here, we demonstrate that endosomal transport of mRNAs encoding mitochondrial proteins orchestrates local bioenergetics in infectious hyphae of *Ustilago maydis*. Using the subunit Atp3 of electron transport chain Complex V as a model, we uncover that the endosomal mRNA transporter Rrm4 is required for efficient mitochondrial protein import, particularly at growth poles. Loss of Rrm4 leads to defects in mitochondrial import, resulting in altered physiology. We propose that endosome-coupled mRNA transport constitutes a fundamental layer of subcellular mitochondrial homeostasis, with implications extending from fungal pathogenicity to neuronal disease.

**Significance Statement:** Fungal pathogens depend on efficient polar growth to execute their infection programs. Consequently, their growing cell poles face a massive local demand of energy, which is supplied by mitochondria. Currently, it is unclear how involved proteins of the mitochondrial electron transport chain (ETC) reach these distant organelles. Here, we combine fungal genetics, metabolomics, transcriptomics and minimal invasive live-cell imaging to resolve this spatial challenge in the corn pathogen *Ustilago maydis*. We discover that long-distance endosomal hitchhiking of mRNAs encoding mitochondrial ETC components is essential to sustain active mitochondria at the expanding pole. Ultimately, this membrane-coupled mRNA trafficking precisely orchestrates subcellular mitochondrial function, disclosing a previously unrecognized Achille’s heel for the development of novel fungicides.

## Introduction

Highly polarized cells, such as fungal hyphae, must coordinate the function of their organelles, such as nuclei, endosomes and mitochondria over long distances to sustain polar growth (Riquelme *et al*, 2018). This intracellular networking is particularly evident in pathogens that depend on efficient hyphal growth as part of their infection strategy (Gow & Hube, 2012; Lanver *et al*, 2017). Mitochondria play a central role by generating ATP through oxidative phosphorylation. This is most commonly achieved by the electron transport chain (ETC) Complexes I, III, and IV, which establish a proton gradient across the inner mitochondrial membrane. The F_O_F_1_ ATP synthase (complex V) uses this gradient to synthesize ATP.

The vast majority of mitochondrial proteins are encoded in the nucleus (Rath *et al*, 2021). To enter the organelle, many mitochondrial proteins contain N-terminal mitochondrial targeting sequences (MTS), which are recognized by specific receptors present on the outer mitochondrial membrane, while other proteins rely on internal targeting signals (Wiedemann & Pfanner, 2017). Evidence is increasing that mitochondrial protein import is enhanced by local translation at the surface of mitochondria (Cohen *et al*, 2024; Vazquez-Carrada *et al*, 2026). In earlier studies, it was found that 80 S ribosomes were present at the mitochondrial outer membrane (Kellems *et al*, 1974; Kellems & Butow, 1972). More recent cryo-electron tomography studies consistently demonstrate the presence of ribosomes clustering externally at sites of local constrictions of the outer and inner mitochondrial membranes (Chang *et al*, 2025; Gold *et al*, 2017). In line with these observations, sophisticated proximity labelling techniques identified actively translating ribosomes in the immediate vicinity of mitochondria (Fazal *et al*, 2019; Luo *et al*, 2025).

It has been proposed that local protein synthesis at mitochondria is important for coordinating the assembly of membrane complexes. Such local translation is promoted by RNA-binding proteins that recognize specific localization motifs, typically found within the 3’ untranslated region of mRNAs (3’ UTR; Béthune *et al*, 2019; Gadir *et al*, 2011). However, an outstanding question is how mRNAs reach mitochondria located far from the nucleus. This is particularly important in highly polarised cells like fungal hyphae and neurons (Müntjes *et al*, 2021; Vazquez-Carrada *et al*., 2026). In neurons, it was recently reported that a distinct subset of axonal mRNAs encoding mitochondrial proteins depends on vesicle mediated transport. Loss of transport causes defects in protein import and mitochondrial activity (De Pace *et al*, 2024).

The eukaryotic microorganism *Ustilago maydis* is a model system for long-distance mRNA transport (Vazquez-Carrada *et al*., 2026; Vollmeister *et al*, 2012). This plant pathogenic fungus is the causative agent of corn smut disease. Essential for infection is a switch from yeast-like proliferation to highly polarised hyphal growth. The resulting infectious hyphae grow with a defined axis of polarity: they expand at the apical pole and insert septa at the basal pole in regular intervals, thereby generating cytoplasm-free compartments (Fig. 1A). During this phase of the life cycle, fungal growth depends intensively on microtubule-dependent transport (Müntjes *et al*., 2021; Vazquez-Carrada *et al*., 2026). As a result, defects in long-distance trafficking lead to aberrant bipolar growth (Fig. 1A).

**Fig. 1.**
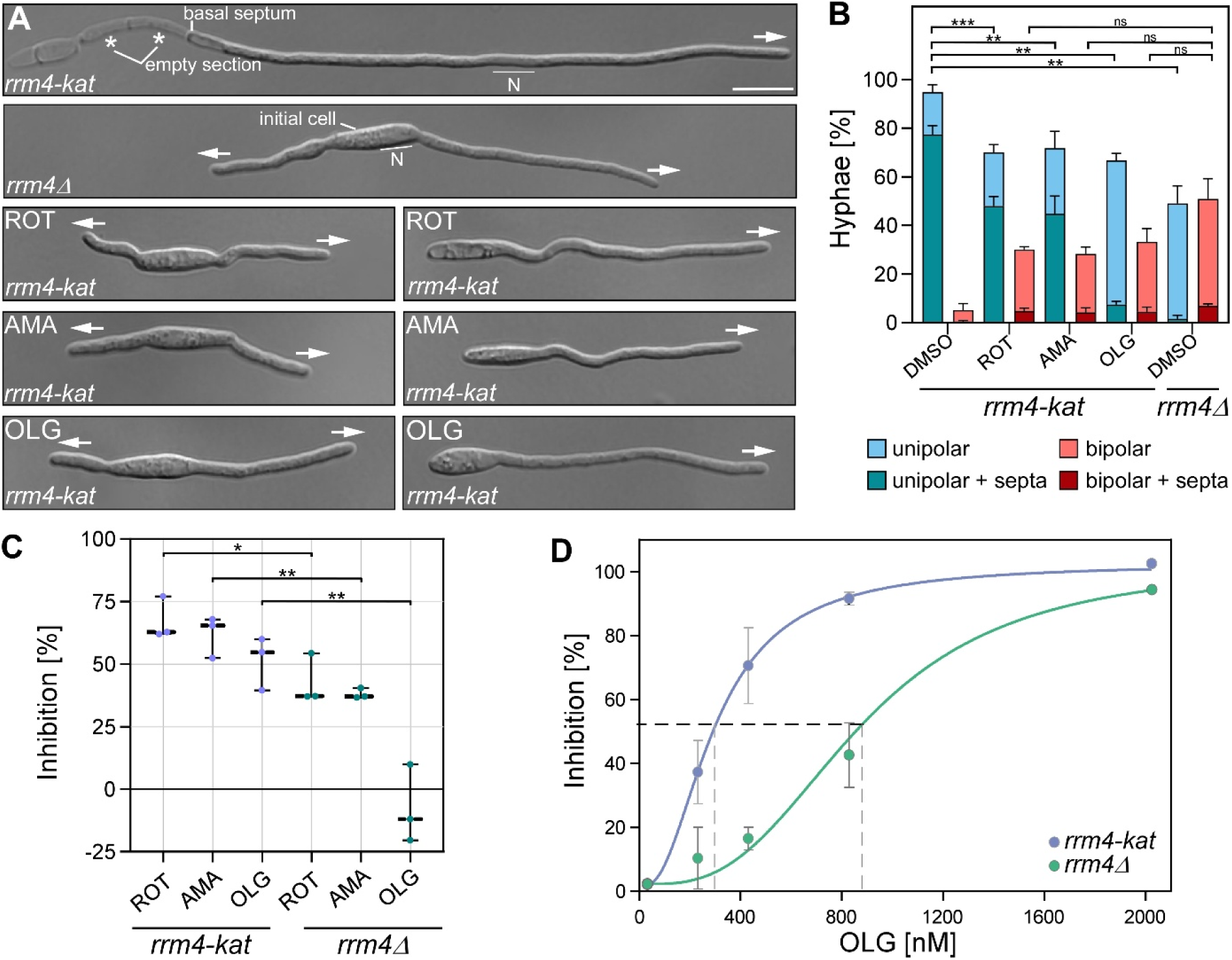
Physiological response to mitochondrial ETC inhibitors. (**A**) Investigation of hyphal morphology upon mitochondrial inhibition with Rotenone (ROT), Antimycin A (AMA), and Oligomycin (OLG) 6 hours post-induction (scale bar: 10 µm, N = nucleus, arrows indicate direction of growth). (**B**) Quantification of hyphal morphology following inhibitor treatment (n=3, *N*>100 hyphae). Statistical significance relative to the untreated control was determined using one-way ANOVA with Dunnett’s post-hoc test. (**C**) Normalized hyphal length as a percentage of DMSO control for wild-type and *rrm4*Δ strains across all mitochondrial inhibitors (n=3, *N*>200 hyphae). Statistical significance was determined using one-way ANOVA with post-hoc Tukey’s test to allow for comparisons between both strains and treatments. (**D**) Determination of the inhibitory concentration (IC_50_) for oligomycin in wild-type and *rrm4*Δ backgrounds (n=3, *N*>30 hyphae). The IC_50_ values were calculated using non-linear regression, and statistical significance between strains was determined using Student’s t-test. Error bars represent SEM; statistical significance is indicated by: ***, p<0.001; **, p<0.01; *, p<0.05; ns, not significant.

Rab5a-positive transport endosomes shuttle along microtubules by the coordinated activity of a plus-end-directed Kinesin-3 type motor and minus-end-directed cytoplasmic dynein (Steinberg, 2012). mRNAs are functionally important cargos of this long-distance transport and are carried in a translationally active state (Baumann *et al*, 2014). Consequently, ribosomes are co-transported by this endosomal hitchhiking mechanism (Baumann *et al*., 2014; Higuchi *et al*, 2014). The key mRNA transporter is the RNA-binding protein Rrm4. It contains three C-terminal MademoiseLLE domains that form a protein-protein interaction platform for endosomal attachment via the endosomal adaptor Upa1 (Devan *et al*, 2022; Devan *et al*, 2024; Pohlmann *et al*, 2015).

Rrm4 contains three RNA recognition motifs (RRM) domains at its N-terminus for binding of cargo mRNAs (Olgeiser *et al*, 2019; Stoffel *et al*, 2026; Stoffel *et al*, 2025). Prominent examples include mRNAs encoding septins, where co-translational transport of septin mRNAs is needed for formation of heteromeric complexes of the translation products on the cytoplasmic surface of endosomes. Trafficking of *de novo*-synthesised heteromeric septin complexes is essential to form gradients of higher-order filaments at the growth pole (Baumann *et al*., 2014; Zander *et al*, 2016). These findings establish endosomal mRNA transport as a platform for spatially coordinated protein synthesis and functional deployment.

Transcriptome-wide analyses using individual-nucleotide resolution UV crosslinking and immunoprecipitation (iCLIP) and iCLIP2 revealed that Rrm4 binds hundreds of mRNAs, suggesting that Rrm4-dependent transport contributes to the distribution of bulk mRNAs throughout the hypha (Olgeiser *et al*., 2019; Stoffel *et al*., 2026; Stoffel *et al*., 2025). A closer inspection of mRNAs with Rrm4 binding sites in the 3’ UTR revealed an enrichment for transcripts encoding mitochondrial ETC subunits (Olgeiser *et al*., 2019). In most cases, these binding sites were detected near stop codons, suggesting a co-translational coupling (Olgeiser *et al*., 2019; Stoffel *et al*., 2026; Stoffel *et al*., 2025). In line with this, an earlier proteomic study of membrane-associated proteins showed altered abundance of mitochondrial proteins including Atp4 and complex I subunit Nuo2 in *rrm4*Δ strains (Koepke *et al*, 2011). Given the central role of mitochondria in cellular metabolism, perturbations in Rrm4-dependent mRNA transport are expected to impact not only mitochondrial protein homeostasis but also cellular bioenergetics and metabolic state. Here, we combine physiological, systems-level and mechanistic analyses to establish a direct link between Rrm4-mediated endosomal mRNA transport and mitochondrial protein import, revealing its central role in subcellular mitochondrial homeostasis in infectious hyphae.

## Results

### Hyphal growth is more resistant to mitochondrial inhibition in the absence of Rrm4

To investigate the physiological importance of mitochondrial ETC function in relation to endosomal transport during hyphal growth, we performed a dose-response inhibitor study using the ETC inhibitors Rotenone (ROT, Complex I), Antimycin A (AMA, Complex III), and Oligomycin (OLG, Complex V; Fig. EV1A-([A-Z])). Inhibitor treatment at the respective IC_50_ (defined as the inhibitor concentration causing 50% inhibition of hyphal length) not only affected hyphal length, but also significantly increased the fraction of aberrant, bipolar cells (Fig. 1A-B; *rrm4-kat* in genetic background of laboratory strain AB33 expressing functional fusion protein of Rrm4 with the fluorescence protein mKate2). Notably, this resembles the aberrant *rrm4*Δ phenotype regarding both the induction of bipolarity and reduced amounts of basal septa (Fig. 1B). Live-cell imaging demonstrated that pharmacological treatment did not abolish ATP-dependent shuttling of Rrm4-Kat-positive endosomes (Fig. EV1F). Hence, hyphae still generate sufficient ATP to support polar growth (Fig. EV1C-([A-Z])).

Comparing control strains with the *rrm4*Δ mutant revealed that in the absence of Rrm4 the reduced hyphal growth caused by ETC inhibitors was less prominent (Fig. 1C; Fig. EV1G). In the case of OLG, growth of *rrm4*Δ hyphae was hardly impaired (Fig. 1C, Fig. EV1G). A more refined analysis of OLG inhibition revealed that the IC_50_ in the *rrm4*Δ background was approximately 3-fold higher than in the control strain (860 nM vs. 280 nM, respectively; Fig. 1D; Fig. EV1B, H), suggesting that loss of Rrm4 improved the ability of hyphae to cope with Complex V inhibition. In essence, inhibiting mitochondrial respiration phenocopies the loss of Rrm4 with respect to hyphal growth, and *rrm4*Δ strains are more resistant to mitochondrial inhibitors. A plausible explanation for this observation is a switch to respiratory-independent ATP production, such as glycolysis.

### Rrm4-deficient hyphae bypass mitochondrial dysfunction through compensatory glycolytic flux

To systematically characterize the cellular impact of Rrm4 deficiency, we conducted a multi-layered omics analysis, beginning with transcriptomic profiling six hours post-induction (h.p.i.) of hyphal growth. Comparing the *rrm4Δ* mutant to the wild-type revealed a distinct transcriptional rewiring, identifying 414 genes with decreased mRNA abundance and 134 genes with increased mRNA abundance (Fig. 2A; Fig. EV2A). KEGG pathway enrichment analysis of these differentially expressed genes highlighted a major shift in cellular priorities. Genes with reduced mRNA levels showed a significant enrichment for steroid biosynthesis as well as the metabolism of storage polymers and sucrose (Fig. 2B, Table S1). Conversely, genes with increased mRNA levels displayed a robust enrichment for core primary metabolic pathways, most prominently carbon metabolism processes such as glycolysis and gluconeogenesis (Fig. 2C).

**Fig. 2.**
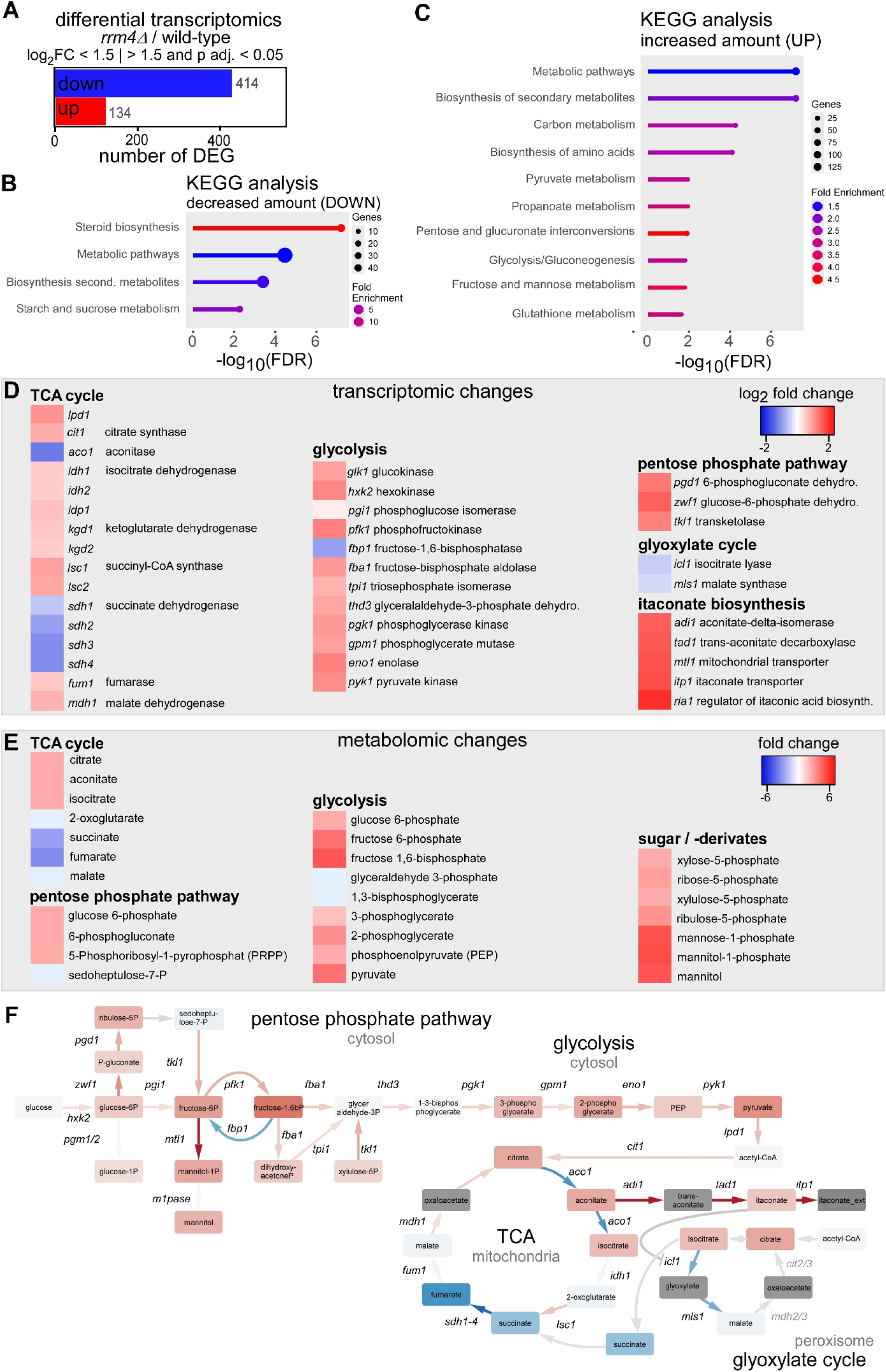
Differential transcriptomic and metabolomic profiling. (**A**) Transcriptomic analysis of *rrm4*Δ vs. wild-type hyphae 6 h.p.i. identifying mRNA targets with significantly increased (red, n=134) and reduced (blue, n=414) abundance (threshold: |*log_2_FC*| > 1.5; *p-adj* < 0.05). (**B-C**) Analysis of KEGG pathway enrichment for genes with differential mRNA abundance. (**D**) Heatmaps showing differential mRNA levels (*log_2_FC*) of genes encoding enzymes for the TCA cycle, glycolysis, pentose phosphate pathway, glyoxylate cycle, and itaconate biosynthesis. (**E**) Heatmap of metabolite ratios (fold-change, FC) comparing mutant and wild-type levels for sugars, glycolysis intermediates, and TCA cycle components. (**F**) Integrated multi-omics network of carbon metabolism mapped across cellular compartments (cytosol, mitochondria, peroxisome). Node and edge colour represent metabolite and transcript *log_2_FC*, respectively. Itaconate directly inhibits isocitrate lyase (Icl1) of the glyoxylate cycle in *Mycobacterium tuberculosis*, indicating a functional link between itaconate biosynthesis pathway and the observed repression of glyoxylate cycle encoding genes (Ruetz *et al*, 2019). Additional genes encoding iso-enzymes of Mdh1 and Cit1 were identified potentially catalyzers in the glyoxylate cycle in peroxisomes. The respective genes are shown in light grey as their transcript-levels did not change significantly.

Prompted by these broad transcriptional changes, we performed a detailed metabolomic investigation to assess the functional biochemical state of the cells (Fig. 2E; Fig. EV2C). Principal component analysis (PCA) of intracellular metabolites confirmed that wild-type and *rrm4Δ* hyphae form two highly distinct metabolic populations (Fig. EV2D). Overall profiling revealed profound alterations in intracellular metabolite pools, characterized by a massive accumulation of sugar derivatives and a severe disruption within mitochondrial metabolic cycles.

Combining these major multi-omics findings reveals a clear, compensatory redirection of central carbon metabolism. Both methods independently demonstrate that *rrm4Δ* cells undergo a massive metabolic shift away from a malfunctioning mitochondrial tricarboxylic acid (TCA) cycle, diverting instead toward cytosolic glycolysis. At the transcriptional level, there was a broad repression of mRNAs encoding key TCA cycle enzymes, specifically aconitase (*Aco1*) and succinate dehydrogenase subunits (Sdh1-4/Complex II; Fig. 2D). This perfectly mirrors a functional bottleneck observed in the metabolomic data, where upstream metabolites like citrate and cis-aconitate strongly accumulate while downstream intermediates like succinate and fumarate are significantly depleted (Fig. 2E). To bypass this mitochondrial dysfunction, the cells apparently accelerate glycolysis. We observed a consistent transcriptional upregulation of key glycolytic enzymes, including phosphofructokinase (*Pfk1*) and pyruvate kinase (*Pyk1*), paired with the downregulation of the gluconeogenic marker fructose-1,6-bisphosphatase (*Fbp1*). Correspondingly, metabolomic data showed a robust accumulation of glycolytic intermediates, directly reflecting this heightened flux. Furthermore, both datasets indicate that excess carbon, unable to pass through the defective TCA cycle, is channeled into alternative overflow pathways. This is evidenced by increased transcripts for alternative respiration (*aox1*) and the itaconate biosynthesis pathway (*adi1*, *tad1*, *itp1*), alongside a consistent reduction in glyoxylate cycle genes (*icl1*, *mls1*).

By mapping these multi-omics datasets to their respective cellular compartments, we generated an integrated metabolic network to visualize this rewiring (Fig. 2F). This comprehensive map clearly contrasts the highly active cytosolic glycolytic pathway—driven by upregulated enzymes and characterized by the accumulation of glucose 6-phosphate and fructose 6-phosphate—with the pronounced bottleneck in the mitochondrial TCA cycle. Within the mitochondria, the network pinpoints the exact site of this blockage, showing citrate accumulating precisely at the nodes where *Aco1* and *Sdh1-4* expression is repressed. Ultimately, this integrated model demonstrates that the loss of Rrm4 triggers a mitochondrial crisis, forcing the cell to adapt by accelerating glycolytic flux and diverting excess organic acids into alternative overflow pathways.

### Loss of Rrm4 causes defects in mitochondrial protein import of *de novo*-synthesized Atp3

Having established that *rrm4*Δ cells undergo a metabolic shift, we sought to understand the underlying molecular trigger. Because these mutants showed specific resistance to Oligomycin (Fig. 1), we hypothesized a malfunctioning of the F_O_F_1_-ATP synthase (Complex V). The bipartite genetic origin of the ATP synthase makes its assembly highly dependent on coordination between nuclear and mitochondrial gene expression (Fig. 3A). Thus, altered endosomal transport of mRNAs encoding nuclear-encoded synthase subunits may provide a mechanistic link between Rrm4 function and complex V activity. Consistently, we found a significant enrichment of binding sites in the 3’ UTRs and/or at stop codons of mRNAs encoding subunits of the complex. These targets include a wide range of subunits such as Atp1 (α), Atp2 (β), and Atp3 (γ), as well as various membrane-associated components like Atp4 (b), Atp7 (d), and Atp14 (Fig. 3A, Table S2; Olgeiser *et al*., 2019; Stoffel *et al*., 2025).

**Fig. 3.**
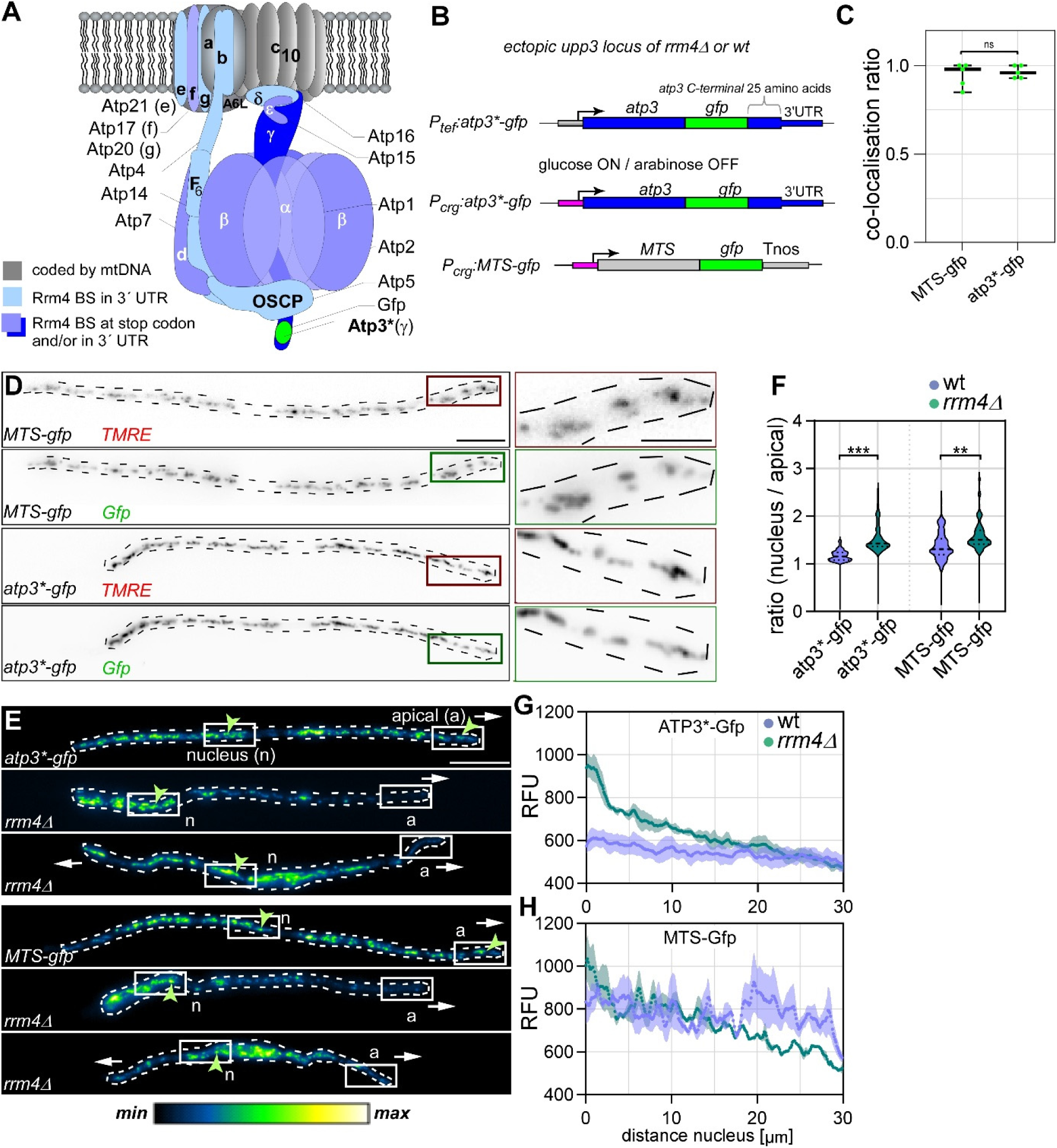
Validation of Atp3*-Gfp as a functional mitochondrial component and its spatial distribution. (**A**) Structural schematic of the mitochondrial F_O_F_1_ ATP synthase complex highlighting subunits encoded by nuclear DNA and mitochondrial DNA (mtDNA). Binding sites of Rrm4 identified by iCLIP are located within the 3’ UTRs and/or the stop codons of nuclear encoded targets (modified from Kucharczyk *et al*, 2009; Olgeiser *et al*., 2019; Stoffel *et al*., 2026). (**B**) Schematic representation of the reporter constructs. All constructs are C-terminal Gfp fusions but are expressed under different promoters and integrated at distinct genomic loci: (1) Atp3 expressed from its native promoter at the *atp3* locus, (2) Atp3 and (3) the corresponding MTS control construct under the control of the *P_crg1_* promoter at the *upp3* locus. The Atp3 “sandwich” fusions (marked as *) are specifically designed to preserve the native Rrm4 binding site environment at the stop codon and within its 3’ UTR. (**C**) Quantification of the co-localization ratio between MTS-Gfp, Atp3*-Gfp and the mitochondrial dye TMRE using Pearson’s correlation coefficient, and (**D**) representative micrographs (size bars 10 µm, and 5 µm in inset) (**E**) Micrographs of Gfp reporter distribution with defined regions of interest near the nucleus (n) and the apical end (a) (A; size bar, 10 µm, white arrows indicate direction of growth, green arrows indicate XY).). (**F**) Comparison of nucleus-to-apical end fluorescence ratios for Atp3*-Gfp and MTS-Gfp in the presence (ON) and absence (OFF) of Rrm4 (n=3 experiments, *N*>100 hyphae). (**G-H**) Fluorescence intensity profiles (RFU) for Atp3*-Gfp and MTS-Gfp as a function of distance from the nucleus (n=3, *N*>30; error bars represent SEM) Statistical significance between groups was determined using Student’s t-test with Benjamini-Hochberg correction. Significance degrees: ***, p<0.001; **, p<0.01; *, p<0.05; ns, not significant; error bars represent SEM.

To investigate the subcellular mitochondrial import of Complex V subunits *in vivo*, we tested the functionality of several candidates after tagging them with Gfp at their C-terminus (Fig. 3B; Fig. EV3A). However, fusion to Atp1, Atp2, or Atp4 critically impaired function of the respective proteins, resulting in short, aberrant hyphae (Fig. EV3A-([A-Z])). In contrast, tagging Atp3 with Gfp resulted in a functional reporter (Fig. 3C-G). Notably, we used a special sandwich configuration to preserve the native Rrm4 binding site environment at the stop codon and in its 3’ UTR (Fig. 3B, designated Atp3*). Quantification of hyphal length confirmed that cells were indistinguishable from wild-type hyphae (Fig. EV3C). Microscopic analysis 6 h.p.i. of hyphal growth showed that Atp3*-Gfp accurately co-localizes with the mitochondrial marker TMRE (tetramethylrhodamine ethyl ester; Fig. 3C). This ratio was statistically identical to the mitochondrial targeting sequence control (MTS-Gfp), confirming the fidelity of the reporter (Fig. 3D). Furthermore, Western-blot analysis confirmed the stable production of the approximately 70 kDa Atp3*-Gfp reporter in both wild-type and *rrm4*Δ backgrounds (Fig. EV3D) confirming Atp3*-Gfp as a suitable functional reporter.

To evaluate the spatial distribution of mitochondrial proteins, we monitored the localization of Atp3*-Gfp and MTS-Gfp reporters expressed from the arabinose-inducible *P_crg1_* promoter (Fig. 3E; Bottin *et al*, 1996). In wild-type hyphae (AB33), Gfp signals for both reporters were distributed evenly throughout the cell, reaching mitochondria at both poles (Fig. 3E-H). In contrast, *rrm4*Δ hyphae displayed a significant gradient with decreasing fluorescence intensity towards the growth pole for both the specific Atp3*-Gfp reporter and the general MTS-Gfp control (Fig. 3G-H). Quantification of the nucleus-to-tip fluorescence ratio confirmed a marked depletion of these proteins at the apex of transport-deficient cells compared to the nuclear region (Fig. 3F). Notably, defects were slightly more pronounced for the Rrm4 target Atp3*-Gfp.

### A minimally invasive system to quantify Rrm4-dependent protein dynamics *in vivo*

To directly assess the role of Rrm4 in mitochondrial protein import, we first examined the localization of Atp3*-Gfp in *rrm4*Δ strains. However, interpretation was confounded by pronounced morphological defects, including more bipolar growing cells with increased cell diameter around the nucleus, as well as by non-physiological transcriptional induction of the reporter (Fig. 4A-B, Fig. EV3H). To decouple the specific role of mRNA transport from these baseline developmental defects, we followed a more sophisticated experimental strategy (Fig. 4).

**Fig. 4.**
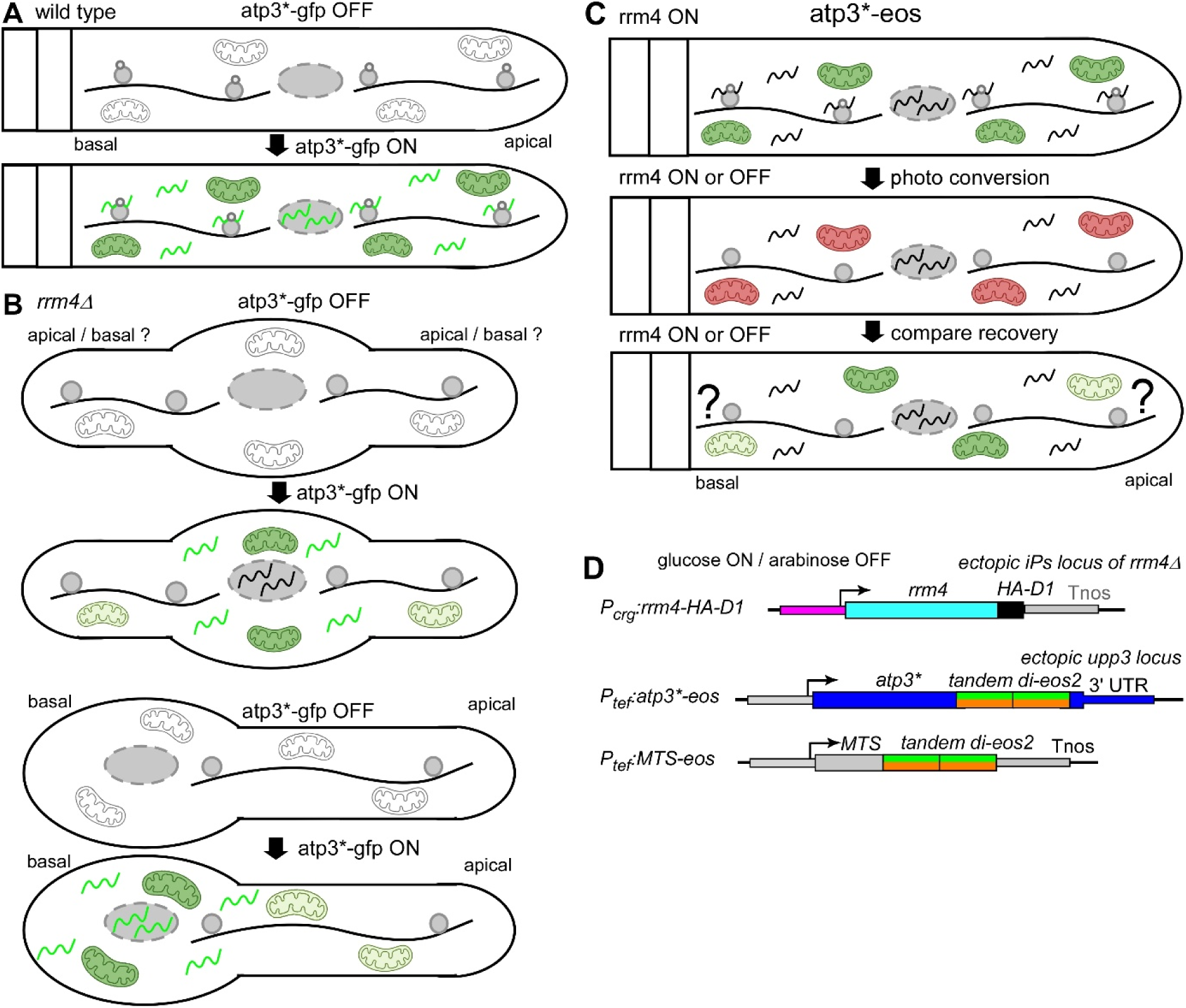
Experimental strategy to assess local mitochondrial protein import. (**A-B**) Schematic models comparing wild-type (**A**) and *rrm4*Δ (**B**) hyphae, highlighting potential biases in protein distribution assessments caused by the aberrant bipolar growth and shortened hyphal phenotypes of the *rrm4*Δ mutant. (**C**) Experimental workflow designed to monitor the spatial Atp3 gradient following the conditional depletion of Rrm4-mediated endosomal transport in established unipolar hyphae. (**D**) Schematic of the constructs used in the bipartite strategy. This includes the conditionally depletable allele (*rrm4-HA-D1*) for rapid clearance of the transporter, and the photoconvertible tdEos2 fluorescent fusions (Atp3*-Eos and MTS-Eos) used to visually separate *de novo*-synthesized proteins from the pre-existing cellular pool.

First, we developed a regulatable version of Rrm4 in order to switch off the endosomal mRNA transport machinery after establishment of unipolar hyphal growth (Fig. 4C-D). To this end, we placed *rrm4* under the control of the arabinose-inducible *P_crg1_* promoter and fused Rrm4 to a D1 degron sequence (Rrm4-HA-D1; Fig. 4D). This dual-layer control allowed for the rapid depletion of Rrm4, effectively halting transport (Fig. EV4 B-D).

Second, to track dynamic mitochondrial protein uptake without invasive transcriptional induction, we employed the photoconvertible protein Eos (tandem dimeric Eos fluorescent protein tdEos2; Fig. EV4A; Zhou *et al*, 2018) fused to either Atp3* or the MTS control as a minimal invasive biosensor for dynamic mitochondrial protein uptake (Fig. 4D). By switching off *rrm4* expression and measuring the recovery of green fluorescence after an initial photoconversion to red, we could selectively detect newly synthesized proteins while minimizing morphology-related differences (Fig. 4C).

Before applying this system to mitochondrial targets, we validated the photoconversion strategy using Rrm4 itself as a benchmark for dynamic transport (Fig. 5A; Fig. EV5A-([A-Z])). Before photoconversion, Rrm4-Eos displayed characteristic shuttling patterns in the green channel (Fig. 5B-C), which were indistinguishable from the established Rrm4-Kat control (Fig. 5D-G; Müntjes *et al*, 2020). Exposure to minimal invasive violet light (405 nm) resulted in a substantial conversion of the green signal to red (80%), effectively depleting the pool of green-fluorescent Rrm4-Eos while maintaining the structural integrity of the transport units (Fig. 5D-E). After four hours of recovery newly synthesized green Rrm4-Eos signals appeared and exhibited shuttling at wild-type velocities (Fig. 5F-G). Importantly, total signal number and velocity remained stable throughout the process (Fig. 5H-I) and the recovery rate reached ∼50% (Fig. 5J, Fig. 5EVF). In essence, the photoconversion strategy reliably captures dynamic Rrm4 transport, thereby establishing a foundation for its application to mitochondrial targets (Fig. 4C).

**Fig. 5.**
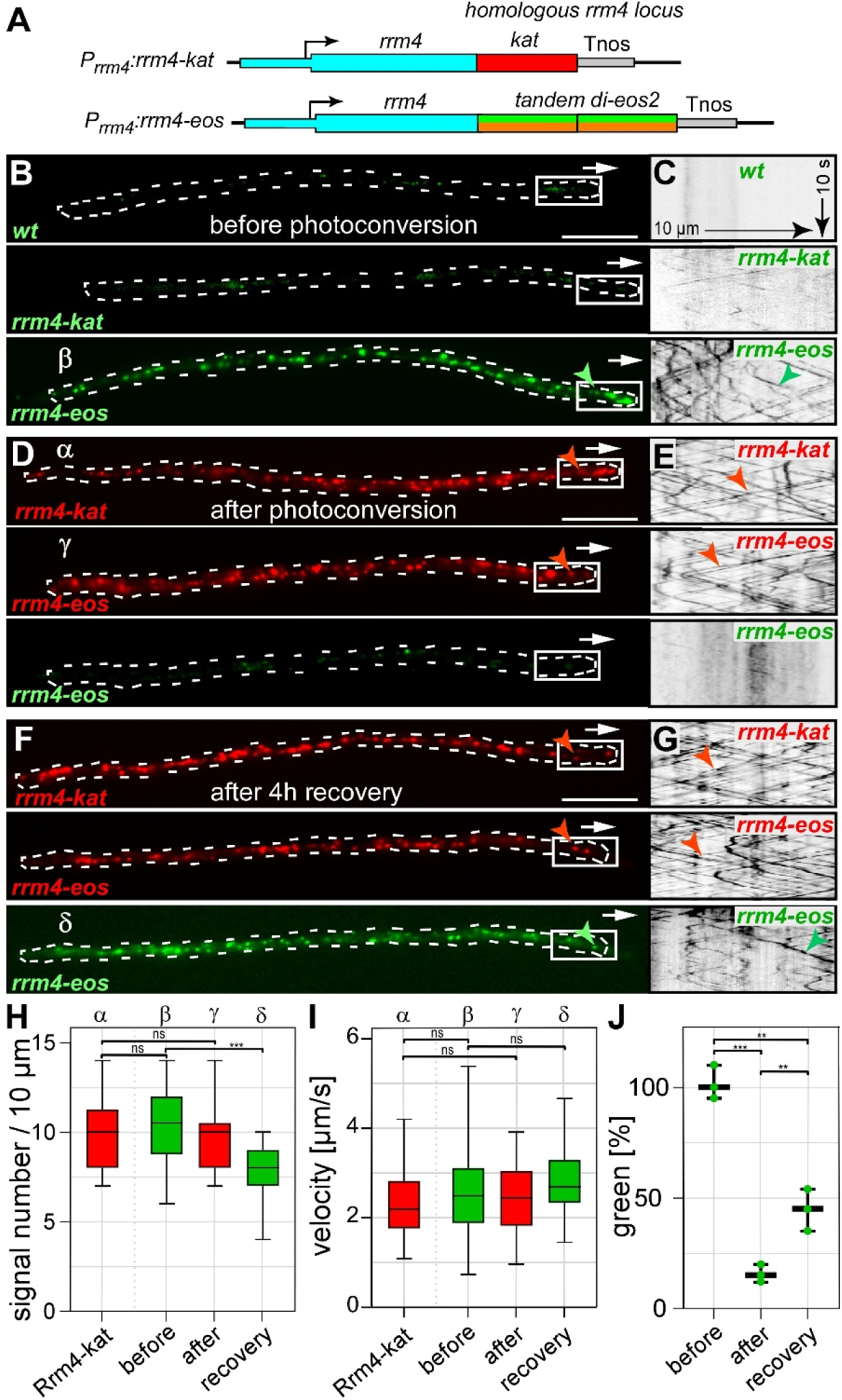
Validation of the Rrm4-Eos photoconversion system. (**A**) Constructs for integration of *rrm4-kat* (control) and *rrm4-eos* at the native genomic locus. (**B-C**) Micrographs and kymographs showing movement of Rrm4-Kat and Rrm4-Eos prior to minimally invasive violet light exposure (405 nm; scale bar, 10 µm, white arrows indicate direction of growth, green arrows indicate XY). (**D-E**) Micrographs and kymographs directly following violet light exposure, illustrating the immediate shift in fluorescence signal. (**F-G**) Micrographs and kymographs showing fluorescence signal and movement 4 h after photoconversion. (**H-I**) Quantification of signal number per 10 μm and particle velocity in μm/s (n=3 experiments, N>30 hyphae). (**J**) Quantification of signal intensity before, immediately after photoconversion and 4 hours later (n=3, *N*>30). Error bars in H, I and J represent SEM. Statistical significance was determined using one-way ANOVA with post-hoc Tukey’s test. Statistical significance is indicated by: ***, p<0.001; **, p<0.01; *, p<0.05; ns, not significant.

### Rrm4-mediated mRNA transport governs localization of Atp3 specifically at apical mitochondria

To study subcellular mitochondrial protein import we employed the functional Atp3*-Eos reporter and the mitochondrial marker control MTS-Eos in unipolar hyphae (Fig. 6A). Atp3*-Eos and the MTS-Eos accurately co-localized with the mitochondrial dye TMRE (Fig. EV6A-([A-Z])) and both recovered efficiently after photoconversion (Fig. 6B). Monitoring the spatial distribution of newly synthesized MTS-Eos in transport-competent hyphae (Rrm4-ON) revealed a relatively uniform distribution (Fig. 6A, C). In contrast, import of newly synthesized Atp3*-Eos was more efficient at apical mitochondria compared to mitochondria at the basal pole near the septum (Fig. 6A, C). Importantly, the distribution of newly synthesized Atp3*-Eos was highly sensitive to the presence of the transport machinery (Fig. 6D). Upon depleting Rrm4, import of newly synthesized Atp3*-Eos was strongly reduced at the apical region, but increased near the basal septum (Fig. 6D). In contrast, mitochondrial localization of *de novo*-synthesized MTS-Gfp was not affected (Fig. 6D).

**Fig. 6.**
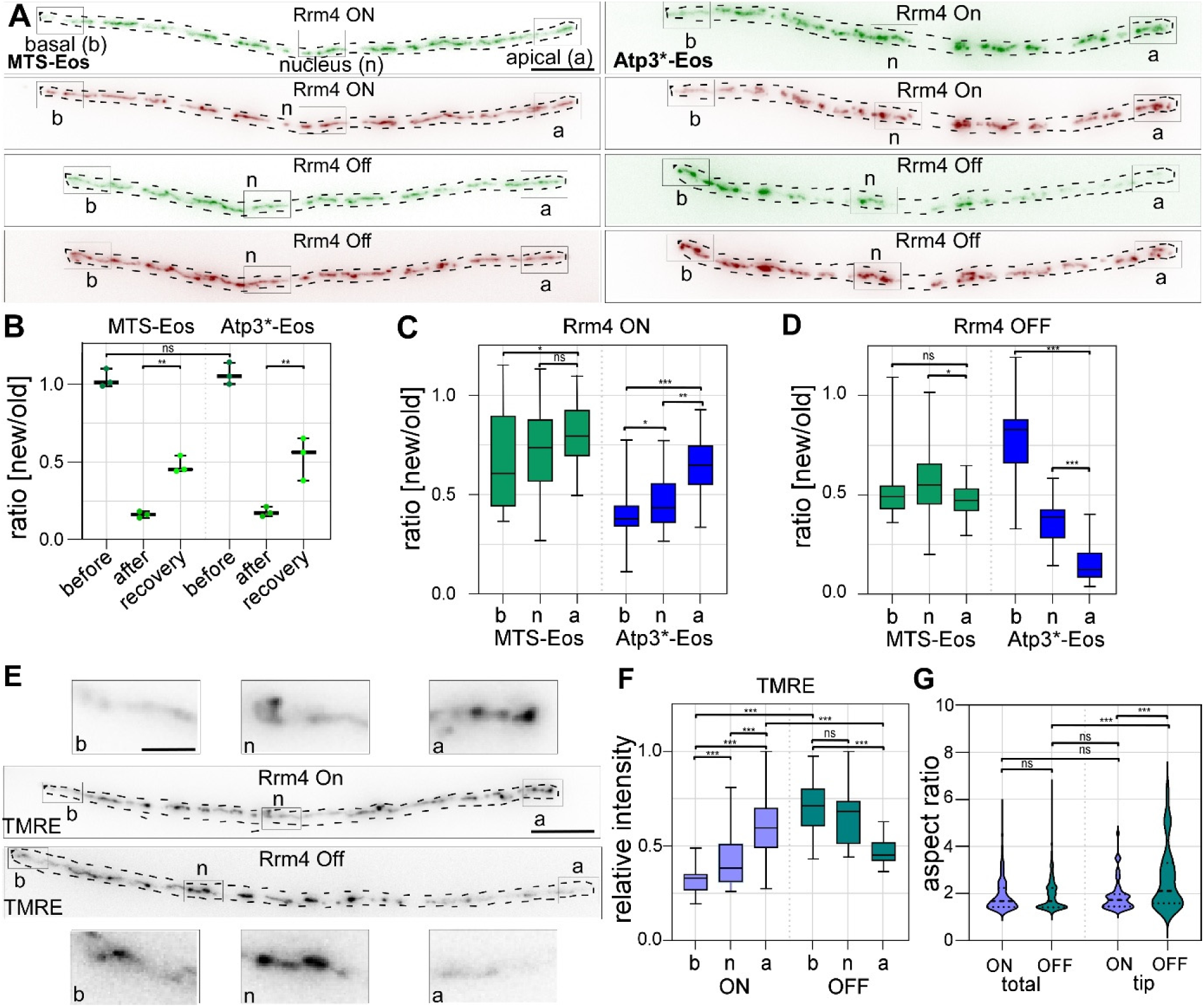
Rrm4-dependent distribution of Atp3 and mitochondrial activity. (**A**) Fluorescence images of MTS-Eos (left) and Atp3*-Eos (right) in hyphae expressing Rrm4 (top) or lacking Rrm4 expression 4 hours post photoconversion (bottom). Regions of interest (ROIs) at the basal pole (b), nucleus (n) and apical pole (a) are indicated. (**B**) Quantification of signal intensity before, immediately after photoconversion and 4 hours later, expressed as the ratio of green to red fluorescence; statistical significance was determined using one-way ANOVA with post-hoc Tukey’s test. (**C-D**) Quantification of signal intensity before, immediately after photoconversion and 4 hours later in defined subcellular ROIs (b, n, and a) under (**C**) Rrm4-ON and (**D**) Rrm4-OFF conditions. MTS-Eos and Atp3*-Eos data are shown in the same graph for comparison (n=3, *N*>30). Statistical significance between ROIs within each group was determined using one-way repeated measures (RM) ANOVA with Tukey’s post-hoc test to account for linked spatial measurements within individual hyphae. (**E-F**) Images and corresponding regional quantification of TMRE fluorescence at the b, n, and a regions. Statistical significance between Rrm4-ON and Rrm4-OFF conditions for each specific ROI was assessed using Student’s t-test with Benjamini-Hochberg correction for multiple comparisons (n=3, N>30). (Scale bar, 10 µm for micrograph and 2 µm for inset; experimental design given in Fig. EV6E). (**G**) Quantification of mitochondrial aspect ratio at the apical end compared to the total population. Statistical significance between Rrm4-ON and OFF conditions was determined using Student’s t-test (n=3, *N*>30). Significance degrees are indicated as: ***, p<0.001; **, p<0.01; *, p<0.05; ns, not significant; error bars represent SEM.

To determine if this protein depletion impacted mitochondrial physiology, we used TMRE staining as a functional readout (Fig. 6E-F). Since TMRE only sequesters into mitochondria with high membrane potential, it serves as a reliable marker for mitochondrial metabolic activity (Zorova *et al*, 2018). In transport-deficient hyphae, we observed a significant reduction in TMRE fluorescence at the hyphal apex (Fig. 6E-F) correlating with the absence of newly synthesized Atp3*-Eos (Fig. 6A). Notably, the loss of active transport led to a localized change in mitochondrial morphology at the distal pole, while mitochondria in proximal regions appeared normal, those at the apical pole shifted toward a significantly more elongated morphology (Fig. 6G). In essence, Rrm4-mediated mRNA transport is specifically required for the efficient localization of Atp3*-Gfp in apical mitochondria, and this delivery mechanism is essential for maintaining the mitochondrial integrity and activity at the growth pole.

## Discussion

In polarized cells, communication between nucleus and distal organelles needs to be coordinated over long distances. This is of particular importance for mitochondria that are distant from the nucleus to support the high energy demand at the growth tip (Müntjes *et al*., 2021; Vázquez-Carrada *et al*, 2025). Here, we present strong evidence supporting the hypothesis that endosomal transport of nuclear mRNAs encoding mitochondrial ETC components is essential for subcellular mitochondrial protein import and hence local mitochondrial homeostasis.

### The endosomal mRNA transporter Rrm4 is crucial for mitochondrial physiology

The vast majority of mitochondrial proteins are encoded by nuclear mRNAs and translated in the cytoplasm. Although MTS is sufficient for mitochondrial targeting, increasing evidence indicates that local translation near mitochondria enhances import efficiency for specific transcript subsets (Cohen *et al*., 2024; Luo *et al*., 2025; Vázquez-Carrada *et al*., 2025). However, a remaining central question is how these mRNAs reach distant mitochondria in highly polar cells.

One possibility is classical adaptor- and motor protein-mediated transport of mRNAs in ribonucleoprotein granules (Amrute-Nayak & Bullock, 2012; Baumann *et al*, 2020; Das *et al*, 2021; Moissoglu *et al*, 2025). Alternatively, mRNAs encoding mitochondrial proteins can be co-transported with mitochondria over long distances (Bauer & Koppers, 2025; Cohen *et al*, 2022; Harbauer *et al*, 2022; Tong *et al*, 2026). Our results support a vesicle-coupled hitchhiking mechanism, a common mRNA trafficking pathway conserved from fungi to neurons. The endosomal system serves as a mobile platform to orchestrate selective local translation, a fundamental strategy to overcome the physical constraints of highly polarized cell growth (Bauer & Koppers, 2025; De Pace *et al*., 2024; Olgeiser *et al*., 2019; Stoffel *et al*., 2026; Vázquez-Carrada *et al*., 2025). In fungi, the key endosomal mRNA transporter Rrm4 recognizes cargo mRNAs encoding ETC components and is required for mitochondrial delivery of the translation products, thereby supporting mitochondrial activity and metabolism (Müntjes *et al*., 2021; Vázquez-Carrada *et al*., 2025).

Accordingly, our pharmacological studies revealed that treatment of hyphae with mitochondrial inhibitors phenocopied aberrant bipolar growth of *rrm4*Δ mutants, suggesting that efficient ETC function is needed for unipolar growth. Combining transcriptomics with metabolomics disclosed a profound reconfiguration of central carbon metabolism in *rrm4*Δ. The observed upregulation of the glycolytic pathway likely serves as a compensatory mechanism to maintain ATP production via substrate-level phosphorylation when mitochondrial ATP synthesis fails. Furthermore, our data identify a critical bottleneck within the mitochondrial TCA cycle. To mitigate this mitochondrial carbon stress, *U. maydis* diverts accumulated acids into overflow pathways such as itaconate biosynthesis (Hartmann *et al*, 2018). Thus, loss of Rrm4 appears to trigger a mitochondrial deficiency in oxidative phosphorylation that the cell attempts to overcome by accelerating glycolytic flux and exporting excess carbon. Similar defects in maintaining TCA cycle intermediates have been reported for ETC mutants in cultured mouse embryonic fibroblast cells (Ryu *et al*, 2024). Furthermore, in both *S. cerevisiae* and higher eukaryotes, mitochondrial or TCA cycle impairment triggers a metabolic shift where cells upregulate glycolytic flux via specialized signalling pathways to maintain energy homeostasis and biosynthetic precursors when oxidative phosphorylation fails (Pfeiffer & Morley, 2014; Zhao *et al*, 2016). In essence, the mRNA transporter Rrm4 and thus endosomal mRNA transport are crucial for ETC function, coordinating mitochondrial homeostasis and global metabolism of growing hyphae.

### The mRNA transporter Rrm4 governs polar entry of mitochondrial proteins

A plausible explanation for the altered metabolic state of *rrm4*Δ hyphae is inefficient entry of mitochondrial proteins. Initially, we addressed this aspect by comparing wild-type and *rrm4*Δ mutants. However, due to the substantial differences in growth modes we established a more sophisticated experimental strategy relying on regulated Rrm4 expression combined with minimal invasive biosensors to quantify mitochondrial protein import. This extra effort was highly rewarding, since we discovered clear differences in subcellular mitochondrial import and homeostasis (Fig. 7). In wild-type hyphae, Atp3 import is more efficient in mitochondria at the growth pole. Switching off *rrm4* expression in unipolar growing hyphae revealed a general reduction of Atp3 levels in apical mitochondria, accompanied by a decreased mitochondrial membrane potential (ΔΨm). This reduction correlated with altered morphology, suggesting impaired mitochondrial activity. Importantly, we observed a clear difference comparing the mitochondrial protein amount of the Rrm4-dependent ETC component Atp3*-Eos with the MTS-Eos control at apical but not at basal mitochondria. Thus, we hypothesize that Rrm4-mediated endosomal mRNA transport is particularly important for subcellular entry of mitochondria at the growing pole (Fig. 7). Notably, our results are consistent with lysosome-associated mRNA transport in axons. Loss of the BLOC-one-related complex (BORC) disturbs the translocation of transcripts encoding mitochondrial ETC components, leading to reduced amounts of ETC proteins. This correlates with reduced membrane potential and axonal degeneration. However, a direct mechanistic connection to dynamic subcellular protein import remains to be demonstrated in this system (De Pace *et al*., 2024). Notably, bioenergy-dependent mitochondrial heterogeneity is also known in cultured mouse cells (Ryu *et al*., 2024).

**Fig. 7.**
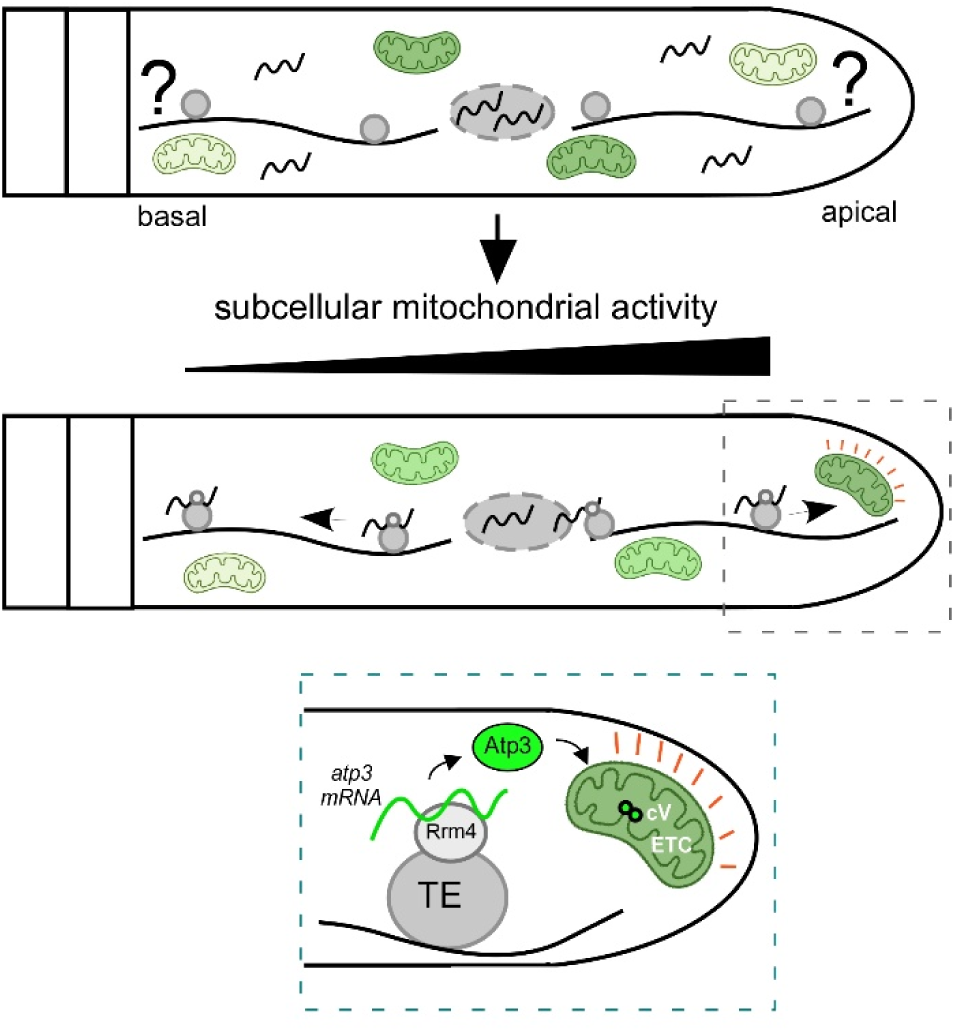
Model depicting how endosomal mRNA transports coordinates subcellular mitochondrial protein import of ETC components. The upper panel illustrates the central biological question: how is the necessary subcellular mitochondrial activity achieved and maintained at distant cellular poles? We hypothesize that Rrm4 recognizes mRNAs encoding subunits of the electron transport chain (ETC), such as *atp3*, for long-distance transport. These transcripts are hitchhiking on transport endosomes (TE) and actively transported along the microtubule cytoskeleton (solid wavy lines) toward the hyphal apex. Upon reaching the growth pole, the mRNAs (green wavy line) are positioned for local translation in the vicinity of mitochondria (green ovals), promoting highly efficient local protein import (Atp3) and ETC assembly (cV, Complex V, small green circles). Consequently, this targeted, Rrm4-dependent mRNA delivery establishes a spatial gradient: subcellular mitochondrial activity progressively increases from the basal pole to the apical pole. This mechanism maintains a high mitochondrial membrane potential and bioenergetic capacity specifically at the apex to support the high energy demands of polar expansion.

The link between localized translation of ETC mRNAs and mitochondrial metabolism has also been studied in non-polar growing cells of *S. cerevisiae* (Tsuboi *et al*, 2020). Mitochondrial association of *atp3* mRNA is enhanced during respiratory metabolism, leading to increased translation that facilitates protein import. In contrast, mRNAs encoding proteins such as the inner mitochondrial membrane component TIM50 exhibited increased mitochondrial association under fermentative conditions, suggesting an intimate link between mitochondrial metabolism, localized translation and protein import (Tsuboi *et al*., 2020).

Rrm4 is required for efficient entry in mitochondria near the growth pole. The underlying mechanism that differentiates between apical and basal mitochondria most likely involves additional RNA-binding proteins. Potential candidates exhibiting specific functions for mitochondrial localization of ETC mRNAs in other systems include the Pumilio family Puf3p from *S. cerevisiae* (Lapointe *et al*, 2018) as well as Cluh1 (Clustered mitochondria homolog Wakim *et al*, 2017) or AKAP1 (outer mitochondrial membrane protein A-kinase anchoring protein 1) from humans (Luo *et al*., 2025). Besides the identification of additional RBPs, it will be crucial to analyze the subcellular site of translation. Endosome-coupled translation has been described for septin mRNAs during microtubule-dependent transport in *U. maydis* (Baumann *et al*., 2014; Zander *et al*., 2016). Alternatively, local translation at the surface of late endosomes is known to promote mitochondrial protein uptake in axons of retinal ganglion cells. Interfering with the function of late endosomes resulted in aberrant length of mitochondria and reduced mitochondrial membrane potential suggesting that endosome-coupled translation is important for mitochondrial import and function (Cioni *et al*, 2019). Alternatively, close contact between the endoplasmic reticulum (ER) and mitochondria at distinct contact sites has been shown to promote mitochondrial protein import using the ER-SURF pathway (ER surface retrieval; Hansen *et al*, 2018; Koch *et al*, 2024). In essence, regulation of localized translation offers a plausible mechanism to govern subcellular mitochondrial physiology.

## Conclusion

Vesicle-mediated hitchhiking of RNAs is a widespread translocation mechanism found in fungi, plants, vertebrates, and humans (Corradi & Baudet, 2020; Corradi *et al*, 2020; Olgeiser *et al*., 2019; Stoffel *et al*., 2026; Vázquez-Carrada *et al*., 2025). Although the precise transport machinery is not identical across species, we demonstrate that the fundamental communication between the nucleus, vesicles, and microtubule-dependent transport of ETC mRNAs is remarkably similar in fungi and neuronal axons. Malfunction of this prominent trafficking route causes significant defects in polar growth of infectious hyphae and triggers axonal degeneration (De Pace *et al*., 2024; Vázquez-Carrada *et al*., 2025), which manifests clinically in severe neurological pathologies (De Pace *et al*, 2026). Ultimately, our findings pave the way to develop novel strategies to fight fungal pathogens and treat human neurodegenerative diseases.

## Materials and methods

### Plasmids, strains, and growth conditions

The *Ustilago maydis* strains used in this study are derivatives of the laboratory strain AB33, in which the genes encoding the bW/bE heterodimer are under the control of the nitrate-inducible *nar1* promoter (Brachmann *et al*, 2001). Plasmids were generated using *Bsa*I-mediated Golden-gate cloning or Gibson Assembly (NEB). Genetic transformations were performed by protoplast-mediated transformation; all integrations (e.g., at the *rrm4*, *ip^s,^* or *upp3* loci) were verified by PCR and Southern blot analysis. Strains generation is described in Supporting Information Table S3; the corresponding plasmids for strain generation are listed in Supporting Information Table S4.

For proliferation, cells were grown at 28 °C in complete medium (CM) supplemented with 1% (w/v) glucose (CM-Glc) or 1% (w/v) arabinose (CM-Ara) and 0.3% (w/v) ammonium sulphate (non-inducing conditions). Hyphal growth was triggered by shifting cells to the inducing nitrate minimal medium (NM) containing 0.3% (w/v) KNO_3_. For standard induction and multi-omics analysis, this medium was supplemented with 1% (w/v) glucose (NM-Glc) or 1% (w/v) arabinose (NM-Ara).

Sensitivity to mitochondrial inhibitors (Rotenone, Antimycin A, Oligomycin) was assessed by adding inhibitors to established hyphae 2 h post-induction (h.p.i.). Hyphal length was quantified after 4 h of treatment. IC_50_ values were calculated using non-linear regression (log[inhibitor] vs. response) in the programme R.

To switch the transport machinery “off” while maintaining the hyphal state, the cells were grown in CM-Ara and shifted to NM-Ara to induce hyphal growth. After 16 h the medium was replaced with inducing nitrate minimal medium supplemented with 1% (w/v) glucose (NM-Glc). In this state, the nitrate source keeps the cells in the hyphal program by maintaining *nar1*-driven b-complex expression, while glucose acts as a potent repressor of the *crg1* promoter to stop *rrm4* expression. The C-terminal D1-degron tag ensured the rapid clearance of existing Rrm4-HA-D1 protein, with depletion within 2 h of the switch confirmed by Western-blot analysis. This approach allowed for the analysis of mitochondrial import in morphologically normal, unipolar filaments in the absence of Rrm4-mediated transport.

### Transcriptomic analysis

RNA isolation and quality control:

Hyphal growth of *U. maydis* was induced for 6 h.p.i. in NM-Glc. Total RNA was extracted from 50 mL hyphal cultures (OD_600_=0.5) using the RNeasy Plant Mini Kit (Qiagen, Hilden, Germany) following the manufacturer’s instructions. Cell pellets were disrupted mechanically using a Mixer Mill MM400 (Retsch, Haan, Germany) at 30 Hz for 5 min at 4 °C in the presence of glass beads, Buffer RLC, and β-mercaptoethanol. Following mechanical disruption, the lysate was passed through a QIAshredder spin column for additional homogenization. The clarified supernatant was mixed with an equal volume of 70% ethanol and loaded onto an RNeasy spin column. All subsequent steps followed the manufacturer’s protocol, including on-column genomic DNA removal using the RNase-Free DNase Set (Qiagen, Hilden, Germany). RNA integrity and quality were verified using a Bioanalyzer RNA Nano Assay.

Library preparation and sequencing:

Next-generation sequencing (NGS) libraries were prepared using the VAHTS Universal V6 RNA-seq Library Prep Kit for Illumina (Vazyme, NR604-01-02). Sequencing was performed on an Illumina HiSeq 3000 platform to produce 151-nt single-end reads. For the hyphal samples, approximately 10 million raw reads were obtained per sample.

Bioinformatic processing and differential expression:

Initial quality control of raw sequencing data was conducted with FastQC. Subsequent RNA-seq analysis was performed on the open-source platform Galaxy. Reads were filtered and processed using Trimmomatic (v0.36), which included a 20-nt trim at the 5’ end, shortening of the 3’ end when Phred scores dropped below 30, and the removal of reads shorter than 20 nt. Processed reads were aligned to the *U. maydis* genome (Ensembl release 51) using STAR (v2.7.2b), allowing up to four mismatches per read and restricting mapping to a single locus. Uniquely mapped reads per gene were quantified using htseq-count (v0.9.1). These raw counts served as input for differential expression analysis using the Bioconductor package DESeq2. Transcripts were classified as differentially expressed if they exhibited an absolute fold change (∣log_2_FC∣) >1.5 after shrinkage and an adjusted P-value <0.05, applying the Benjamini-Hochberg correction.

### Metabolite measurements

For metabolite measurements, 100 mL of cell culture was quickly filtered through ice-cold 0.9% NaCl using a vacuum filtration system (Merck Millipore) and collected on nylon membranes (25 mm, 0.45 µm). Membranes were immediately placed into 2 mL tubes preloaded with steel and glass beads (Retsch) and snap-frozen in liquid nitrogen; the entire procedure was completed within 45 s.

Metabolite profiling was adapted from previous protocols (Rabinowitz & Kimball, 2007). Extraction was performed using 1 mL of acetonitrile/methanol/0.1 M formic acid (40:40:20, v/v) supplemented with 2 µM thio-ATP as an internal standard. Cells were disrupted in a bead mill (Retsch; 30 Hz, 2 min) with pre-chilled holders to ensure efficient lysis. Following neutralization with 90 µL of 15% (w/v) ammonium hydroxide and incubation at −20 °C for 30 min, samples were centrifuged (16,000 × g, 10 min, 4 °C). The clarified supernatant was diluted with 5 mL ice-cold water and lyophilized for 48 h (Alpha 1–2 LDplus; CHRIST).

Dried samples were reconstituted in 200 µL water and analysed on a Dionex ICS-6000 ion chromatography system coupled to a Q Exactive Plus quadrupole–Orbitrap mass spectrometer (Thermo Scientific; Schwaiger *et al*, 2017). Anion-exchange separation was performed on a Dionex IonPac AS11-HC column (2 mm × 250 mm, 4 µm) with an AG11-HC guard column at 30 °C. A KOH gradient (5 mM to 85 mM over 35 min) was delivered at 380 µL min⁻¹.

Negative electrospray ionization (ESI) used a methanol-based makeup flow (10 mM acetic acid) at 150 µL min⁻¹. Source settings included a spray voltage of −2.8 kV and a capillary temperature of 300 °C. Full-scan mass spectra (*m/z* 60–800) were acquired at 140,000 resolution. Targeted data processing was performed using Skyline (version 25.1), and peak areas were normalized to internal standards and OD_750_ values recorded at sampling.

Metabolic network reconstruction and multi-omics integration were performed using the *U. maydis* genome-scale model (Liebal *et al*, 2022) and visualized in Cytoscape (v3.10).

### Microscopy and photoconversion assays

Laser-based epifluorescence microscopy was conducted using a Zeiss Axio Observer.Z1, following a previous report (Devan *et al*., 2024). To assess uni- and bipolar hyphal growth, cells were cultured in 20 mL volumes until reaching an OD_600_ of 0.5, after which hyphal growth was induced. After 6 h, more than 150 hyphae per strain were examined for growth behaviour (n = 3). Cells were scored for unipolar and bipolar growth, and for the formation of a basal septum. For the analysis of signal number, velocity, and distance travelled by fluorescence-labelled Rrm4-Eos variants, movies were recorded with an exposure time of 150 ms and 150 frames. Over 20 hyphae were analysed per strain (n = 3). All movies and images were processed and analysed using Metamorph software (version 7.7.0.0, Molecular Devices, Seattle, IL, USA; Müntjes *et al*., 2020). For both micrographs and kymographs, a 20 µm segment from the hyphal tip was analysed. To statistically quantify the signal number and velocity, processive signals covering a distance of more than 5 µm were manually counted. Fluorescence imaging was performed with an attached Visitron laser scanning system using 488 nm excitation with a 525/50 nm emission filter and 561 nm excitation with a 600/50 nm emission filter. To visualize mitochondrial membrane potential, hyphae were stained with 100 nM Tetramethylrhodamine ethyl ester (TMRE). Fluorescence images were acquired as z-stacks using identical microscope settings for all samples (laser power and exposure time). Z-stacks were collected with a step size of 0.2 µm and processed using Fiji/ImageJ software. Maximum-intensity projections were generated from the complete z-stacks for visualization and comparative analysis of mitochondrial morphology. For TMRE-based measurements, all image acquisition and processing parameters were kept constant between conditions to allow comparison of relative fluorescence intensities. Control experiments in wild-type cells confirmed that the switch from NM-Ara to NM-Glc—and the associated change in carbon source - did not inherently alter the mitochondrial membrane potential or its distribution along the hyphal axis (Fig. EV6E, F; Crowley *et al*, 2016). Photoconversion of tdEos2-tagged proteins was achieved by exposing cells to 405 nm LED light for 20 min (Lenard *et al*, 2013). To track *de novo* protein import, the recovery of the green (newly synthesized) signal was quantified 4 h after the existing pool was converted to red.

### Image processing and quantitative analysis

All image processing and secondary data analyses were performed using the open-source software Fiji/ImageJ (version 1.54f or higher). For visualization only, lookup tables (LUTs) were adjusted using the BioVoxxel Toolbox plugin without modification of the underlying raw data. For mitochondrial quantification, background subtraction was applied (rolling ball radius = 50 pixels), followed by segmentation using edge-detection algorithms. Measurements were restricted to defined subapical regions: the septum region (10 µm distal from the basal septum), the nucleus region (10 µm proximal to the nucleus), and the tip region (10 µm proximal to the hyphal apex). For the quantification of mitochondrial protein distribution (Fig.3 G-H), line profiles of fluorescence intensity were recorded along the longitudinal axis of the hyphae, starting from the nucleus towards the hyphal tip. To account for varying hyphal lengths, the resulting raw fluorescence units (RFU) were plotted as a function of distance or normalized to the total cell length.

For the analysis of mitochondrial morphology (Fig. 6G), the Feret diameter (the maximum distance between any two points along the selection boundary) was determined for individual mitochondria. The aspect ratio was calculated by dividing the maximum by the minimum Feret diameter, serving as a robust metric for organelle elongation. Mitochondria were categorized based on their subcellular localization, specifically comparing those in the apical region (tip-ROI) to the total mitochondrial population (total-ROI; Leonard *et al*, 2015). Co-localization was assessed by overlaying fluorescence intensity profiles of the respective channels and quantified using Pearson’s correlation coefficient calculated in Fiji (ImageJ; Dunn *et al*, 2011). For Fig. 6C-D, the recovery of newly synthesized green tdEos2 signal was normalized to the pre-existing red signal within defined ROIs (<u>b</u>asal septum, <u>n</u>ucleus, <u>a</u>pical pole) to generate the new/old signal ratio.

### Western blot analysis

Proteins were extracted via trichloroacetic acid (TCA) precipitation. Samples were resolved by SDS-PAGE (8–12% gels) and transferred to nitrocellulose membranes. Primary antibodies: anti-HA (Sigma, 1:1,000) and anti-GFP (Roche, 1:2,000). Secondary antibodies: HRP-conjugated anti-mouse or anti-rabbit (1:5,000). Signals were visualized using ECL and a ChemiDoc imaging system (Bio-Rad). Protein depletion kinetics were analysed by collecting samples at 0, 1, 2, 3, 4, and 5 h after switching from NM-Ara to NM-Glc.

## Data availability statement

In accordance with FAIR (findable, accessible, interoperable, and reusable) data management principles, the metabolomics and transcriptomics datasets generated in this study, along with their associated metadata and computational workflows, have been compiled into an Annotated Research Context (ARC). The ARC is deposited and securely stored in the DataPLANT DataHUB infrastructure.

## Acknowledgements

We thank Jessica Müller, Drs Carl Haag, Eva Maleckova, Lilli Bismar and Thorsten Langner for their contributions to this work. We acknowledge Drs. Florian Altegoer and Kerstin Schipper for helpful discussions and critical reading of the manuscript, and Dr. Sabrina Zander for assistance with the ARC data repository. We are grateful to Simone Esch and Ute Gengenbacher for excellent technical assistance, and to all members of the Feldbrügge group for helpful discussions. This research was generously supported by grants from the DFG under Germany’s Excellence Strategy EXC-2048/1 - project ID 39068111 to MF and PW, DFG-SFB1535 - project ID 458090666 to MF, NW and PW (project A03, A05 and Z02, respectively). Furthermore, the Graduate School Molecules of Infection MOI-V provided funding to MF and JP. The funders had no role in study design, data collection and analysis, decision to publish, or preparation of the manuscript.

## Expanded view figures

**Fig. EV1.**
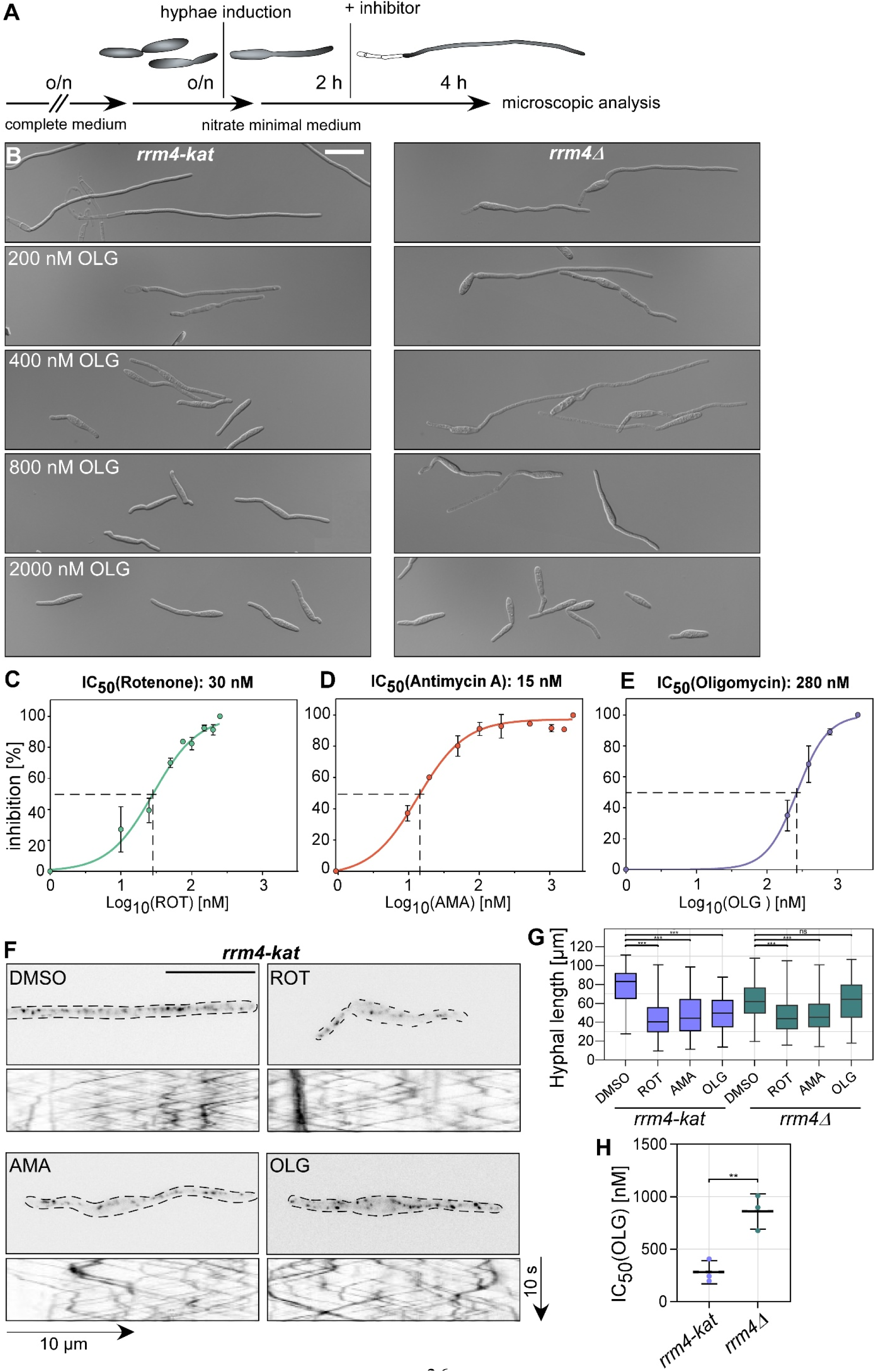
Impact of mitochondrial inhibition on Rrm4 dynamics and growth. (**A–B**) Experimental design and microscopy examples for inhibitor treatments (scale bar, 10 µm). (**C–E**) Investigation of hyphal length to establish IC_50_ values for the indicated inhibitors (n=3 experiments, N>100 hyphae). Dose-response curves were fitted using non-linear regression analysis. (**F**) Micrographs and kymographs of Rrm4-Kat shuttling following treatment with mitochondrial inhibitors (time and distance as indicated by scale bars). (**G**) Comparative growth analysis in wild-type and *rrm4*Δ strains following treatment with ROT (Rotenone), AMA (Antimycin A), and OLG (Oligomycin). Statistical significance relative to the untreated control for each strain was assessed via Student’s t-test with Benjamini-Hochberg correction for multiple comparisons (n=3, *N*>100). (**H**) Comparison of the IC_50_ values specifically for oligomycin in wild-type and *rrm4*Δ backgrounds. Statistical significance was determined using unpaired Student’s t-test, and is indicated by: ***, p<0.001; **, p<0.01; *, p<0.05; ns, not significant; error bars represent SEM (n=3, *N*>30).

**Fig. EV2.**
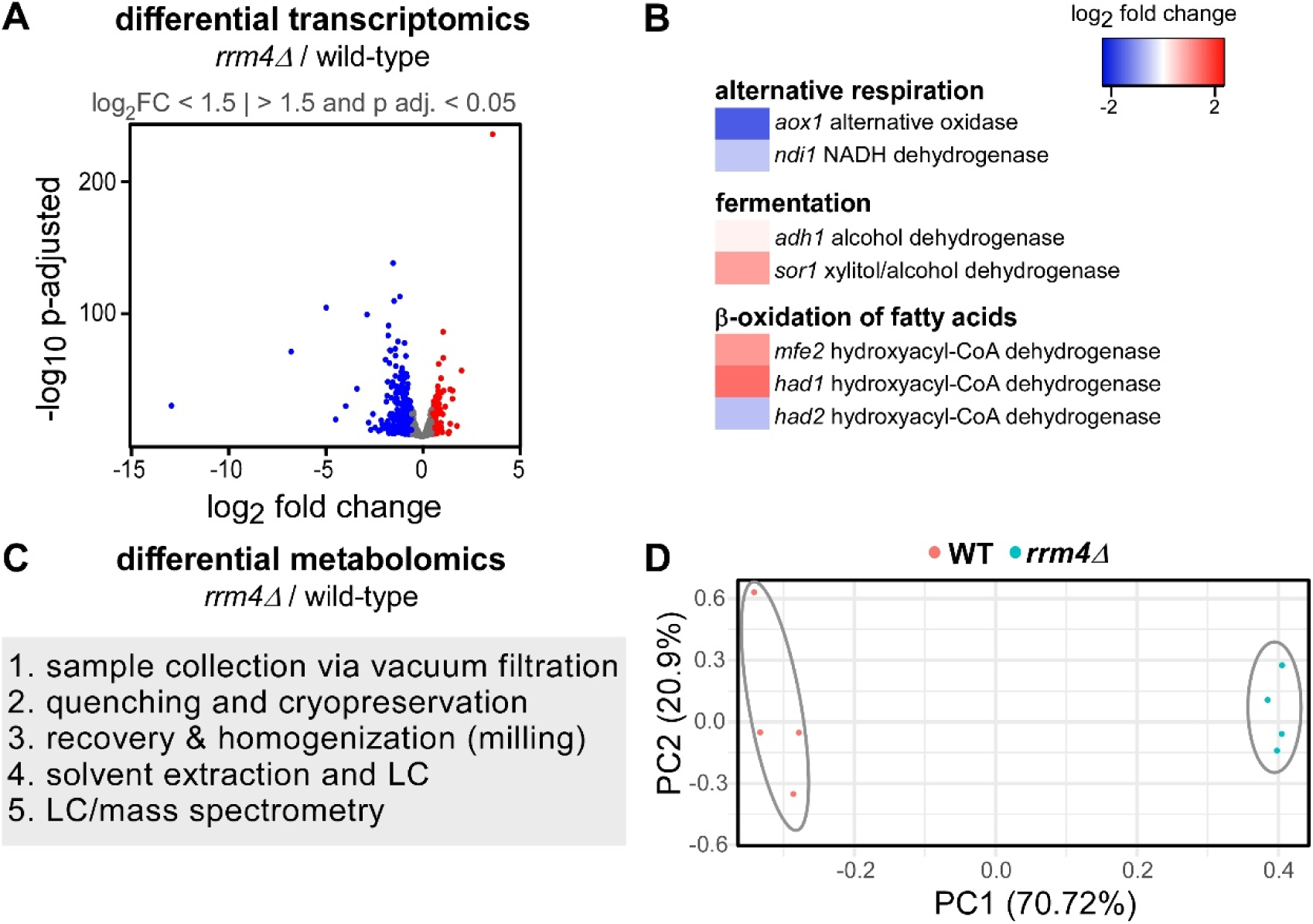
Comparative analysis of wild-type versus *rrm4* mutant. (A) Volcano plot of the transcriptomic comparison between wild-type and *rrm4*Δ mutant cells. (B) Heat map of genes categorized under alternative respiration, fermentation, and β-oxidation. (C) Schematic overview of the sequential steps of sample collection and metabolomic analysis workflow. (**D**) PCA plot of metabolomic profiles for wild-type AB33 and *rrm4*Δ mutant hyphae.

**Fig. EV3.**
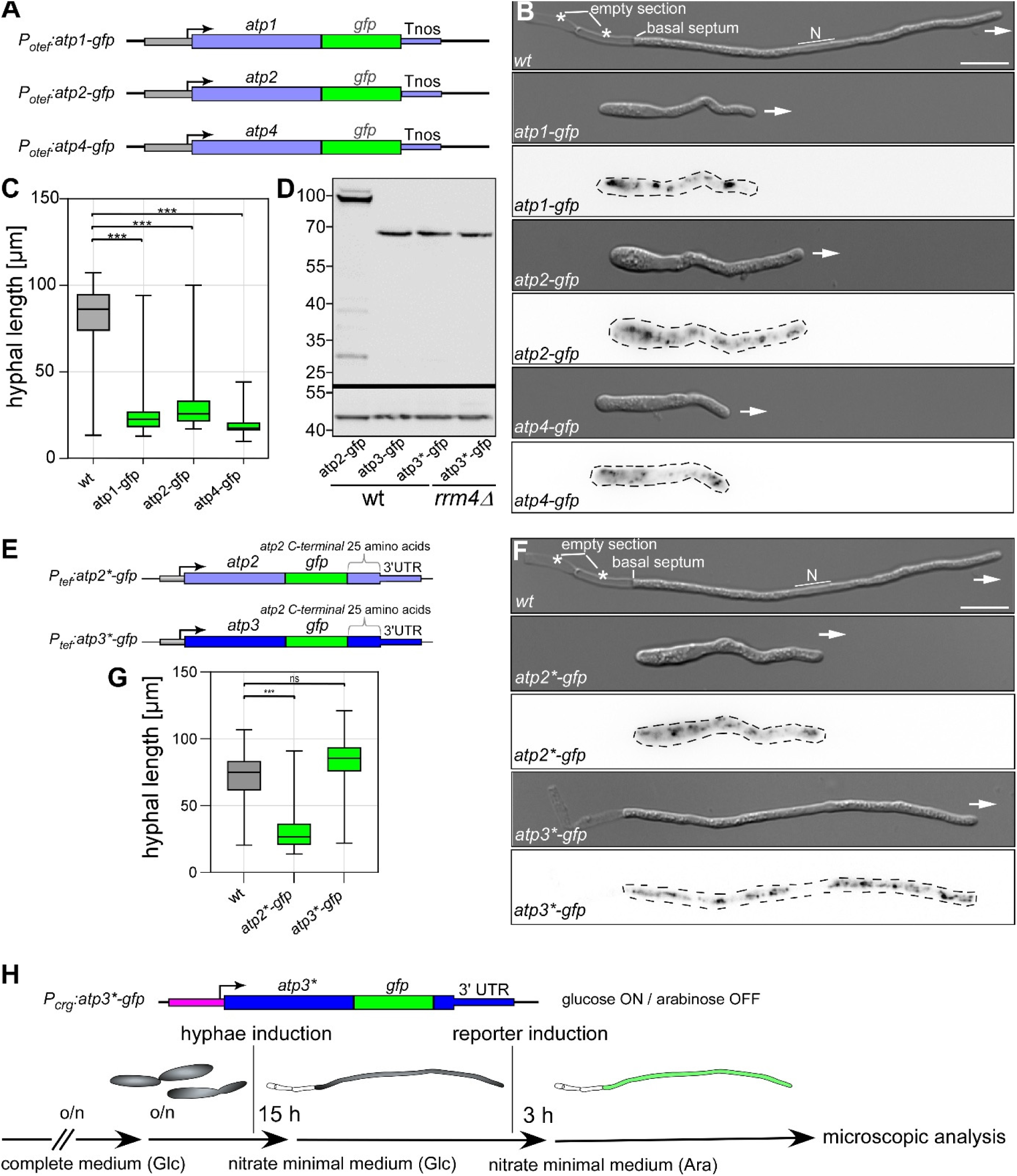
Characterization of reporter functionality of Complex V subunits. (**A**) Schematic representation of the reporter constructs, indicating the insertion sites of the sequence for the Gfp tag. (**B**) Micrographs illustrating the morphological phenotype of strains expressing Atp1-Gfp, Atp2-Gfp, and Atp4-Gfp (scale bar, 10 µm). (**C**) Quantification of hyphal length for the indicated strains. Statistical significance across all groups was determined using one-way ANOVA with Dunnett’s post-hoc test to compare reporters against wild-type. (**D**) Western blot analysis confirming the expression levels of the tested reporters in wild-type and *rrm4*Δ backgrounds. (**E**) Schematic representation of reporter constructs using the constitutively active *P_tef_* promoter and C-terminal “sandwich” fusions marked as *. (**F**) Representative micrographs of hyphae expressing Atp2*-Gfp or Atp3*-Gfp reporters compared to wild-type (scale bar, 10 µm). (**G**) Quantification of hyphal length (n=3, *N*>100; error bars represent SEM and statistical significance was determined using one-way ANOVA with Dunnett’s post-hoc test and is indicated by: ***, p<0.001; **, p<0.01; *, p<0.05; ns, not significant). (**H**) Graphical representation of the experimental workflow used in Fig.3E-([A-Z]) to analyze the spatial distribution of the Atp3*-Gfp reporter. Cells are grown overnight in glucose-containing complete medium, followed by a shift to nitrate minimal medium supplemented with glucose (NM-Glc) for 6 hours to induce hyphal growth while keeping the reporter repressed. To initiate reporter expression, the medium is switched to nitrate minimal medium containing arabinose (NM-Ara). This switch induces the expression of the Atp3*-Gfp reporter under the control of the regulatable *P_crg1_* promoter for 3 hours prior to microscopic analysis.

**Fig. EV4.**
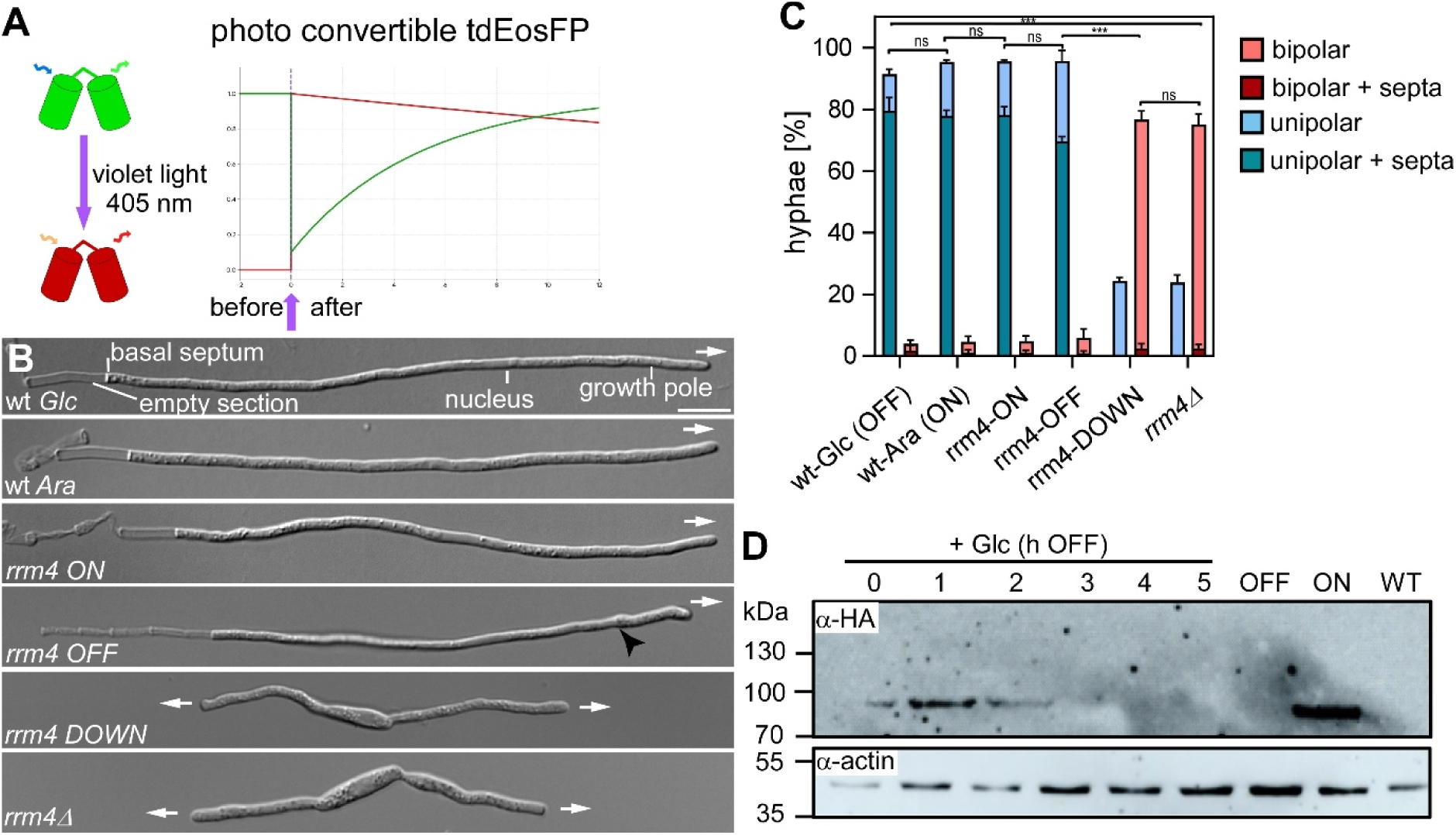
Experimental design for spatiotemporal profiling using photoconvertible tdEos2. (**A**) Schematic of the tdEos2 photoconversion system. Exposure to violet light (405 nm) triggers the irreversible conversion of the fluorophore from a green-emitting to a red-emitting state. The accompanying graph illustrates the theoretical kinetics of fluorescence signal over a 12-hour period: the pre-existing pool is converted to red (red line), while the appearance of newly synthesized protein is tracked via the recovery of the green signal (green line). (**B-C**) Representative DIC micrographs and quantification showing that the Rrm4-HA-D1 strain phenocopies the *rrm4*Δ deletion mutant specifically under repressive conditions (NM-Glc; OFF), where the population shifts from unipolar to bipolar growth. Statistical significance across all groups was determined using one-way ANOVA with post-hoc Tukey’s test (scale bar, 10 µm; n=3, *N*>100). (**D**) Western blot analysis using anti-HA antibody showing the kinetics of Rrm4-HA-D1 depletion over a 5-hour time course after switching to glucose-containing medium; Rrm4-ON (arabinose) and WT (wild-type) serve as controls for expression levels and antibody specificity, respectively. Significance degrees: ***, p<0.001; **, p<0.01; *, p<0.05; ns, not significant; error bars represent SEM.

**Fig. EV5.**
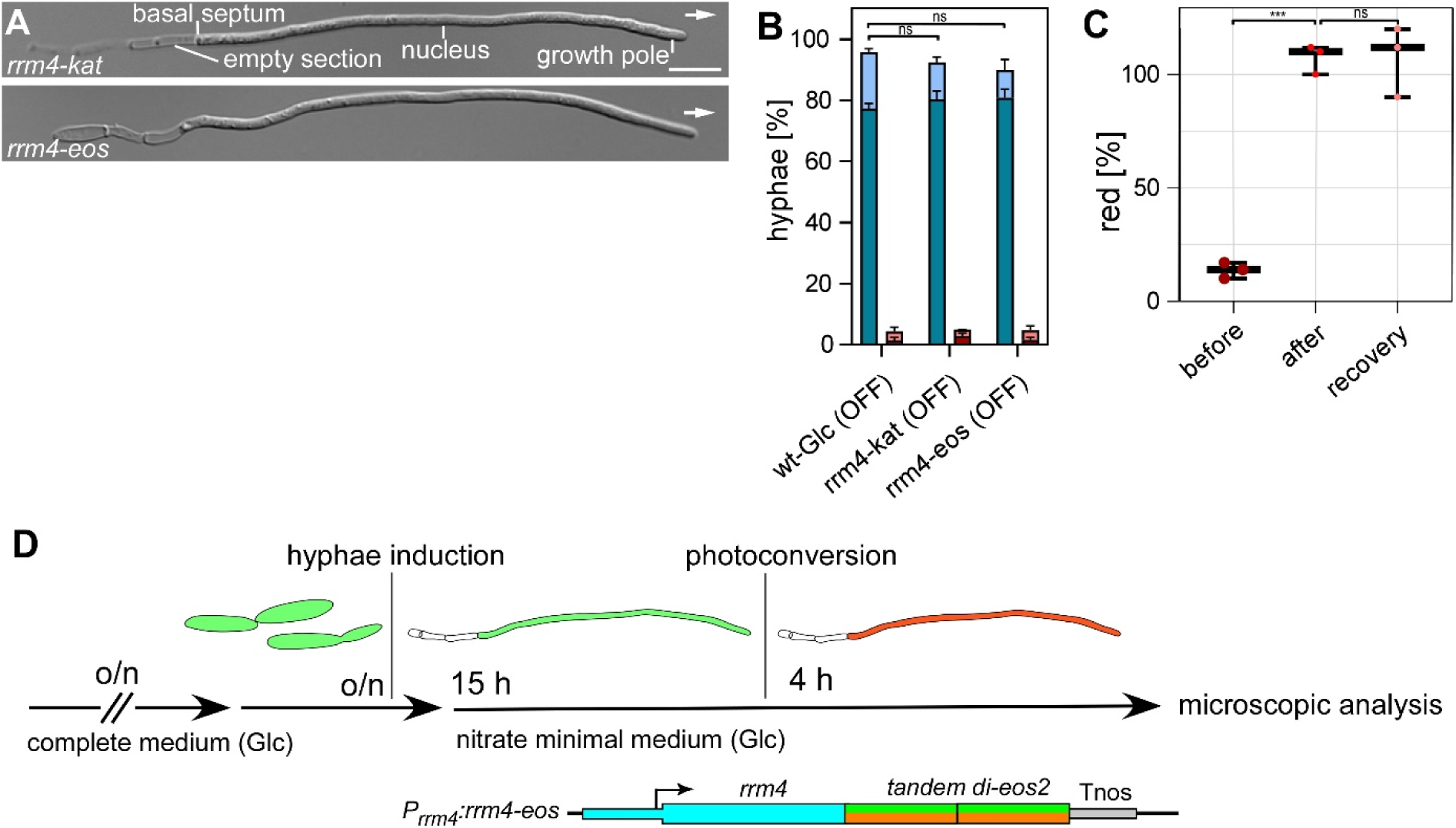
Functional analysis of the regulatable Rrm4-HA-D1 system. (**A**) Representative DIC micrographs showing that hyphal morphology of strains expressing native Rrm4-Kat (rrm4-kat) or Rrm4-Eos (rrm4-eos) is indistinguishable from wild-type (scale bar, 10 µm). (**B**) Quantification of hyphal morphology for strains expressing native Rrm4 fusions after switch from arabinose to glucose, verifying that the carbon source switch does not interfere with unipolar hyphal growth; statistical significance relative to wild-type was determined using one-way ANOVA with Dunnett’s post-hoc test. (**C**) Quantification of red fluorescence Rrm4-Eos signal 4 hours after photoconversion, demonstrating the stability of the photoconverted pool over the recovery period. Statistical significance between time points was determined using one-way ANOVA with post-hoc Tukey’s test (n=3, *N*>30). (**D**) Graphical representation of the experimental workflow for Rrm4-tdEos2 stability assays. Cells are grown overnight in glucose-containing complete medium, shifted to nitrate minimal medium (Glc) to induce hyphal growth while maintaining Rrm4-tdEos2 expression, and subjected to photoconversion followed by microscopic analysis after a 4-hour recovery period. Statistical significance is indicated by: ***, p<0.001; **, p<0.01; *, p<0.05; ns, not significant; error bars represent SEM.

**Fig. EV6.**
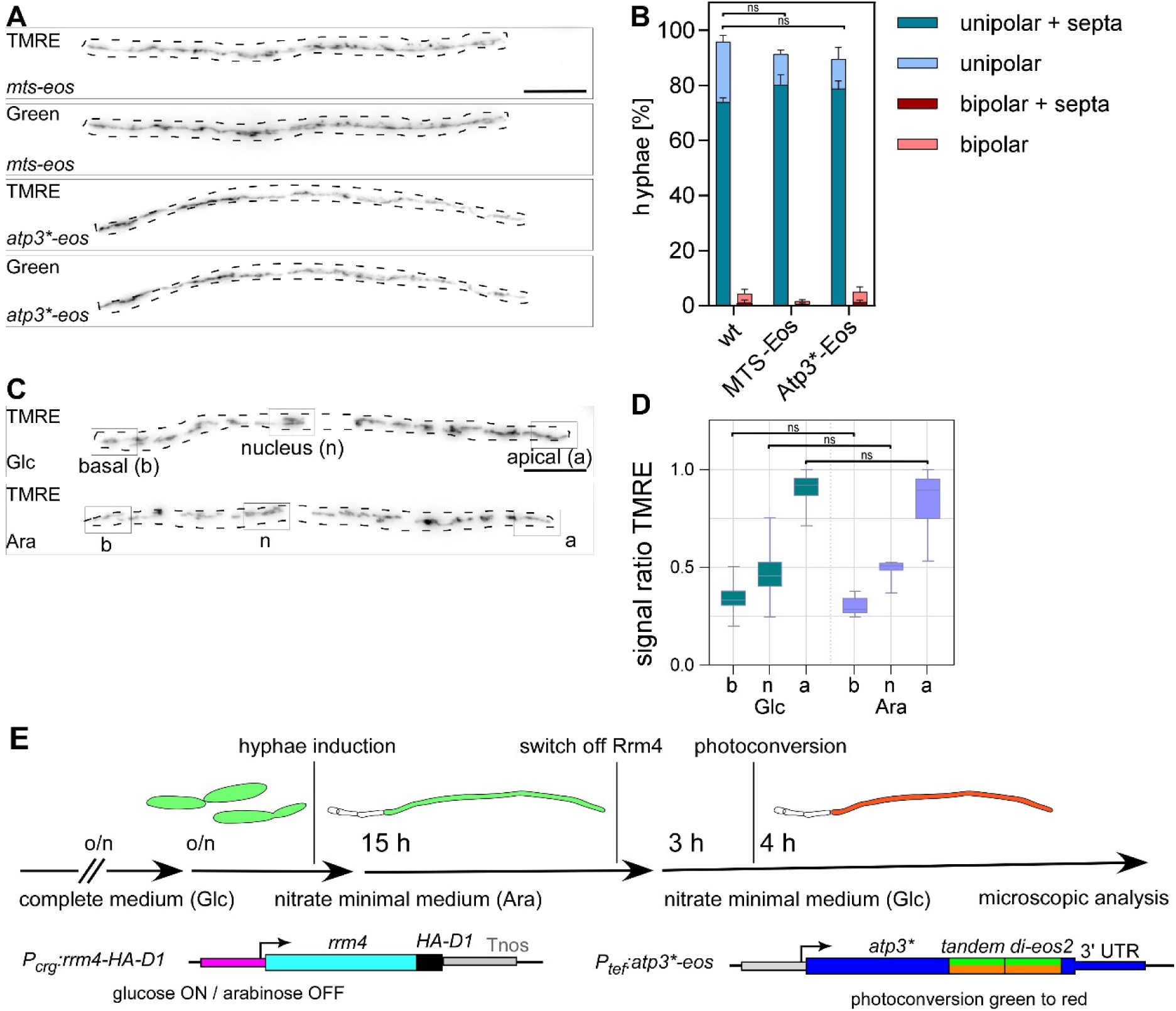
Control and validation of mitochondrial activity reporters. (**A**) Representative micrographs showing co-localization of Atp3*-Eos and MTS-Eos with TMRE in wild-type hyphae (scale bar, 10 µm). (**B**) Morphology and growth analysis confirming that the expression of Eos-tagged reporters does not interfere with hyphal growth. Statistical significance across all groups was determined using one-way ANOVA with post-hoc Tukey’s test (n=3, *N*>100). (**C**) Fluorescence micrographs of wild-type AB33 cells stained with TMRE after growth in either glucose (Glc, repressive) or arabinose (Ara, inductive) conditions to assess the impact of the carbon source on the mitochondrial membrane potential. (**D**) Quantification of the TMRE signal intensity ratios along the hyphal axis, in defined ROIs: the apical pole (a), the nucleus (n), and the basal pole (b). This comparison determines if the carbon source shift independently alters the spatial distribution of mitochondrial activity. Statistical significance between Glc and Ara conditions for each specific ROI was assessed using Student’s t-test with Benjamini-Hochberg correction (n=3, *N*>30). Statistical significance is indicated by: ***, p<0.001; **, p<0.01; *, p<0.05; ns, not significant; error bars represent SEM. (**E**) depicts the graphical representation of the experimental workflow used in Strategy II (results section) to dissect transport-dependent mitochondrial import in unipolar hyphae. Cells are initially grown overnight in complete medium containing arabinose to maintain expression of the regulatable Rrm4-HA-D1 transporter, followed by a 2-hour induction of hyphal growth in nitrate minimal medium (NM-Ara). To achieve the Rrm4-OFF condition, the medium is switched to nitrate minimal medium supplemented with glucose (NM-Glc) to repress the *P_crg1_* promoter. The C-terminal D1-degron tag facilitates rapid clearance of the remaining Rrm4 protein, ensuring full depletion before further analysis. Following this depletion phase, the established unipolar hyphae are subjected to photoconversion, and the recovery of newly synthesized mitochondrial reporters (Atp3*-tdEos2 or MTS-tdEos2) is quantified 4 hours later to assess the impact of abolished endosomal transport.

