## Supplementary Information SI Table 1-4 for "Endosomal mRNA transport coordinates local mitochondrial bioenergetics during polar fungal growth"

$ shared first authorship

* shared corresponding authorship

[1] Heinrich Heine University Düsseldorf, Institute of Microbiology, Cluster of Excellence on Plant Sciences, 40204 Düsseldorf, Germany
[2] Forschungszentrum Jülich GmbH, Institute of Bio- and Geosciences Biotechnology (IBG-1), 52428 Jülich, Germany
[3] Theodor Boveri Institute, Biocenter, University of Würzburg, Am Hubland, 97074 Würzburg, Germany
[4] Heinrich Heine University Düsseldorf, CEPLAS metabolism & metabolomics laboratory, Cluster of Excellence on Plant Sciences, 40204 Düsseldorf, Germany

**Dr. Michael Feldbrügge**

Institute of Microbiology,

Cluster of Excellence on Plant Sciences, Collaborative Research Centre MibiNet

Heinrich Heine University Düsseldorf, 40204 Düsseldorf, Germany

**Table S1. KEGG analysis**

| Grp. | Adj.Pval | Fold | Pathway |
|---|---|---|---|
| **Upregulated** | 3.77E-07 | 1.8 | Metabolic pathways |
| | 2.10E-03 | 2.2 | Biosynthesis of secondary metabolites |
| | 2.23E-03 | 3.4 | Biosynthesis of cofactors |
| | 4.01E-03 | 7.9 | Beta-Alanine metabolism |
| | 8.24E-03 | 6.5 | Glutathione metabolism |
| | 8.24E-03 | 8.6 | Sulfur metabolism |
| | 8.97E-03 | 6 | Arginine and proline metabolism |
| | 2.11E-02 | 6.3 | Pantothenate and CoA biosynthesis |
| | 2.76E-02 | 5.7 | Propanoate metabolism |
| | 3.56E-02 | 7.5 | Pentose and glucuronate interconversions |
| | 4.98E-02 | 4.6 | Glycerophospholipid metabolism |
| | 5.99E-02 | 3.5 | Purine metabolism |
| | 6.12E-02 | 4.1 | Glycine serine and threonine metabolism |
| | 6.12E-02 | 10 | Taurine and hypotaurine metabolism |
| | 7.29E-02 | 3.8 | Cysteine and methionine metabolism |
| | 7.29E-02 | 2.5 | Carbon metabolism |
| | 7.41E-02 | 3.6 | Valine leucine and isoleucine degradation |
| | 7.66E-02 | 7.5 | Lysine biosynthesis |
| | 7.66E-02 | 4.5 | Lysine degradation |
| | 7.66E-02 | 3.5 | Pyruvate metabolism |
| | 9.37E-02 | 4.1 | Glycerolipid metabolism |
| **Downregulated** | 5.91E-06 | 14.4 | Steroid biosynthesis |
| | 9.23E-04 | 1.6 | Metabolic pathways |
| | 1.02E-02 | 2 | Biosynthesis of secondary metabolites |
| | 8.04E-02 | 4.2 | Starch and sucrose metabolism |

**Table S2. Rrm4 target mRNAs encoding Complex V components**

| Category | Subunit | Protein | U. maydis Identifier |
|---|---|---|---|
| F1 | α | Atp1 | UMAG_10213 |
| | β | Atp2 | UMAG_10397 |
| | γ | Atp3 | UMAG_05090 |
| | δ | Atp16 | UMAG_01103 |
| | ϵ | Atp15 | UMAG_10754 |
| | OSCP | Atp5 | UMAG_06324 |
| FO | b | Atp4 | UMAG_10548 |
| | d | Atp7 | UMAG_12050 |
| | e | Atp21 | UMAG_10374 |
| | f | Atp17 | UMAG_10180 |
| | g | Atp20 | UMAG_00975 |
| Accessory Subunits | | Atp18 | UMAG_11576 |
| | | Atp14 | UMAG_02360 |
| | | Inh1 | UMAG_02361 |
| | | Atp10 | UMAG_12212 |
| | | Ap11 | UMAG_03867 |
| | | Atp12 | UMAG_10822 |

### Table S3. Description of *U. maydis* strains used in this study

| Strain name with lab_id | Locus | Progenitor strain | Short description |
|---|---|---|---|
| AB33<br>(UMa133) | *b* | FB2 | *Pnar:bW2bE1*, expression of active b heterodimer under control of the *nar1* promoter, strain grows filamentous upon changing the nitrogen source. (Brachmann et al., 2001) |
| AB33rrm4Δ(UMa273) | *rrm4* | AB33 | carrying a deletion of *rrm4*. (Becht et al., 2005) |
| AB33rrm4-kat(UMa1985) | *rrm4* | AB33 | expressing Rrm4 C-terminally fused to mKate2. (Müntjes et al., 2020) |
| AB33rrm4-eos(UMa1954) | *rrm4* | AB33 | expressing Rrm4 C-terminally fused to tdEos2. (This Study, plasmid pUMa2949) |
| AB33MTS-egfp(UMa1113) | *iPs* | AB33 | expressing MTS mitochondrial signal peptide C-terminal fused to eGfp. (This Study, plasmid pUMa1606) |
| AB33Pcrg:MTS-gfp(UMa1224) | *iPs* | AB33 | expressing MTS mitochondrial signal peptide C-terminal fused to eGfp under control of Arabinose inducible promotor *crg1*. (This Study, plasmid pUMa1604) |
| AB33 rrm4Δ/Pcrg:MTS-egfp(UMa1229) | *iPs* | AB33rrm4Δ | expressing MTS mitochondrial signal peptide C-terminal fused to eGfp under control of Arabinose inducible promotor *crg1*. In the background of *rrm4* deletion. (This Study, plasmid pUMa1604) |
| AB33atp1-gfp(UMa2996) | *upp3* | AB33upp3Δ | expressing Atp1 C-terminal fused to eGfp in *upp3* locus. (This Study, plasmid pUMa3932) |
| AB33atp2-gfp(UMa3051) | *upp3* | AB33upp3Δ | expressing Atp2 C-terminal fused to eGfp in *upp3* locus. (This Study, plasmid pUMa3936) |
| AB33atp4-gfp(UMa2997) | *upp3* | AB33upp3Δ | expressing Atp4 C-terminal fused to eGfp in *upp3* locus. (This Study, plasmid pUMa4176) |
| AB33atp3-gfp-swatp3_3UTR atp3(UL0043) | *upp3* | AB33upp3Δ | expressing Atp3 with an insertion of eGfp 25 AS before C-terminus, fused to 3' untranslated region of atp3. (This Study, plasmid pUMa2369) |
| AB33atp2-gfp-swatp2_3UTR atp2(UL0001) | *upp3* | AB33upp3Δ | expressing Atp2 with an insertion of eGfp 25 AS before C-terminus, fused to 3' untranslated region of atp2. (This Study, plasmid pUL0001) |
| AB33Pcrg:atp3-gfp-swatp3_3UTR atp3(UL0104) | *upp3* | AB33upp3Δ | expressing Atp3 with an insertion of eGfp 25 AS before C-terminus, fused to 3' untranslated region of atp3 under control of Arabinose inducible promotor *crg1*. (This Study, plasmid pUL0084) |
| AB33Pcrg:atp3-gfp-swatp3_3UTR atp3/ rrm4Δ (UL0107) | *upp3*<br>*rrm4* | AB33upp3Δ/ Pcrg:atp3-gfp-swatp3_3UTR atp3 | expressing Atp3 with an insertion of eGfp 25 AS before C-terminus, fused to 3' untranslated region of atp3 under control of Arabinose inducible promotor *crg1*. In the background of Rrm4 deletion. (This Study, plasmid pUL0084) |
| AB33rrm4Δ/Ptef:atp3-eos-swatp3_3UTR atp3/ Pcrg:Rrm4-HA-d1_Tnos (UL0210) | *upp3*<br>*rrm4*<br>*iPs* | AB33rrm4Δ/Pcrg:Rrm4-HA-d1_Tnos | expressing Atp3 with an insertion of tdEos2 25 AS before C-terminus, fused to 3' untranslated region of atp3. In the background of Rrm4 under control of Arabinose inducible promotor *crg1* fused to degron sequence d1, for degradation after turn off. (This Study, plasmid pUL0342) |
| AB33rrm4Δ/Ptef:MTS-eos / Pcrg:Rrm4-HA-d1_Tnos (UL0216) | *upp3*<br>*rrm4*<br>*iPs* | AB33rrm4Δ/Pcrg:Rrm4-HA-d1_Tnos | expressing MTS mitochondrial signal peptide C-terminal fused to tdEos2. In the background of Rrm4 under control of Arabinose inducible promotor *crg1* fused to degron sequence d1, for degradation after turn off. (This Study, plasmid pUL0346) |
| AB33rrm4Δ/Pcrg:Rrm4-HA-d1_Tnos(UL0038) | *rrm4*<br>*iPs* | AB33rrm4Δ | Rrm4 under control of Arabinose inducible promotor *crg1* fused to degron sequence d1, for degradation after turn off in the background of *rrm4* deletion. (This Study, plasmid pUMa4770) |

**Table S4. Description of plasmids generated for *U. maydis* strain generation**

| Plasmid | Plasmid ID | Resistance cassette | Short description |
|---|---|---|---|
| rrm4-eos_Tnos_natR | pUMa2949 | natR | Integrative plasmid for the generation of an rrm4-tdEos2 fusion gene at the native *rrm4* locus of *U. maydis*. In this construct, the stop codon of rrm4 is replaced by a eos gene. Transcription is terminated by the nos terminator. Nourseothricin resistance cassette is used as the selection marker. |
| MTS-egfp_Tnos_narR | pUMa1606 | natR | Integrative plasmid for the generation of a MTS signal sequence fused to gfp gene at the *iPs* locus of *U. maydis*. Nourseothricin resistance cassette is used as the selection marker. |
| Pcrg:MTS-egfp_Tnos_narR | pUMa1604 | natR | Integrative plasmid for the generation of a MTS signal sequence fused to gfp gene under control of the arabinose *pcrg1* promotor, at the *iPs* locus of *U. maydis*. Nourseothricin resistance cassette is used as the selection marker. |
| rrm4Δ_natR | pUMa1750 | natR | Integrative plasmid for knockout of the rrm4 gene in *U. maydis*. The *rrm4* gene is replaced by a nourseothricin resistance cassette (Becht et al., 2005). |
| Potef_atp2-egfp-atp2_Tnos_natR | pUMa3936 | natR | Integrative plasmid for the generation of an atp2-egfp fusion gene at the *upp3* locus of *U. maydis*. Transcription is terminated by the nos terminator. Nourseothricin resistance cassette is used as the selection marker. |
| Potef_atp1-egfp-atp2_Tnos_natR | pUMa3932 | natR | Integrative plasmid for the generation of an atp1-egfp fusion gene at the *upp3* locus of *U. maydis*. Transcription is terminated by the nos terminator. Nourseothricin resistance cassette is used as the selection marker. |
| Potef_atp4-egfp-atp2_Tnos_natR | pUMa4176 | natR | Integrative plasmid for the generation of an atp4-egfp fusion gene at the *upp3* locus of *U. maydis*. Transcription is terminated by the nos terminator. Nourseothricin resistance cassette is used as the selection marker. |
| atp2-gfp-swatp2_3UTR atp2 | UL0001 | natR | Integrative plasmid for the generation of an atp2 with an insertion of eGfp 25 AS before C-terminus, fused to 3' untranslated region of atp2, at the *upp3* locus of *U. maydis*. Transcription is terminated by the nos terminator. Nourseothricin resistance cassette is used as the selection marker. |
| Atp3-gfp-swatp3_3UTR atp2 | UL0043 | natR | Integrative plasmid for the generation of an atp3 with an insertion of eGfp 25 AS before C-terminus, fused to 3' untranslated region of atp3, at the *upp3* locus of *U. maydis*. Transcription is terminated by the nos terminator. Nourseothricin resistance cassette is used as the selection marker. |
| Pcrg:atp3-gfp-swatp3_3UTR atp3 | UL0084 | natR | Integrative plasmid for the generation of an atp3 with an insertion of eGfp 25 AS before C-terminus, fused to 3' untranslated region of atp3, at the *upp3* locus of *U. maydis*. Transcription is terminated by the nos terminator. Nourseothricin resistance cassette is used as the selection marker. |
| Pcrg:Rrm4-HA-d1_Tnos | pUMa4770 | cbxR | Integrative plasmid for the generation of Rrm4 under control of Arabinose inducible promotor *crg1* fused to degron sequence d1, for degradation after turn off in the background in *iPs* locus for cbx resistance. |
| Atp3-eos-swatp3_3UTR atp3 | pUL0342 | natR | Integrative plasmid for the generation of atp3 with an insertion of tdEos2 25 AS before C-terminus, fused to 3' untranslated region of atp3, at the *upp3* locus of *U. maydis*. Transcription is terminated by the nos terminator. Nourseothricin resistance cassette is used as the selection marker. |
| Ptef:MTS-eos | pUL0346 | natR | Integrative plasmid for the generation of MTS mitochondrial signal peptide C-terminal fused to tdEos2, at the *upp3* locus of *U. maydis*. Transcription is terminated by the nos terminator. Nourseothricin resistance cassette is used as the selection marker. |
